# Dual regulation of chemical stress-induced *DDI2/3* expression by a transcription factor Fzf1 and nucleosome in *Saccharomyces cerevisiae*

**DOI:** 10.64898/2026.05.06.723303

**Authors:** Ying Du, Aiyang Lin, Joshua A.R. Brown, LeAnn J. Howe, Wei Xiao

## Abstract

*DDI2* and *DDI3* (*DDI2/3*) are duplicated genes in *Saccharomyces cerevisiae* that exhibit strong induction by a transcription factor Fzf1 in response to chemical treatments like cyanamide (CY) and methyl methanesulfonate (MMS). Although, like *DDI2/*3, *SSU1*, *YHB1* and *YNR064C* also contain an Fzf1-binding consensus sequence CS2 and are coordinately regulated by Fzf1, these genes are only modestly induced by CY and MMS. To identify additional *cis*-acting elements in the *DDI2/3* promoter, we made *DDI2/3* promoter deletions in a reporter system and identified upstream repressing sequences (URS) spanning 480 nucleotides. To test a hypothesis that the chromatin structure constitutes the URS, we utilized a yeast strain capable of histone H3/H4 depletion by shifting carbon sources. Following histone depletion, *DDI2/3* were strongly induced in an Fzf1 dependent manner, while *YHB1* was repressed. Interestingly, under histone depletion conditions, CY or MMS treatment further increased expression of all Fzf1-regulated genes to comparable levels in an Fzf1 dependent manner. A genome-wide MNase-seq analysis showed that CY treatment reduced the nucleosome occupancy at the mapped *DDI2/3* URS region in wild-type cells, but not in in *fzf1*Δ cells. These findings collectively indicate that Fzf1 plays dual roles in regulating the *DDI2/3* response to CY. Firstly, it binds CS2 and serves as a transcription activator. Secondly, it is required for the chromatin remodeling at URS. This two-tier regulation at the *DDI2/3* promoter helps to explain why *DDI2/3* achieve much higher fold induction by CY and MMS than other Fzf1-regulated genes, suggesting Fzf1 to be a candidate pioneer transcription factor.

**Impact statement:** A budding yeast transcription factor Fzf1 can regulate a group of genes in response to chemical stresses through two independent mechanisms, namely *cis-trans* promoter binding and chromosomal remodeling.

## Introduction

*Saccharomyces cerevisiae DDI2* and *DDI3* (*DDI2/3*) are duplicated genes located on chromosomes VI and XIV, respectively. They share nearly identical coding and promoter sequences with only a few nucleotide sequence variations (Supplementary Fig. S1). These genes were strongly induced by a DNA-damaging agent methyl methanesulfonate (MMS) (Fu et al., 2008; Gasch et al., 2000; Jelinsky and Samson, 1999). *DDI2/3* is homologous to a gene in the soil fungus *Myrothecium verrucaria* that encodes cyanamide (CY) hydratase (Maier-Greiner et al., 1991), are robustly induced by CY (Li et al., 2015), and Ddi2/3 indeed possess cyanamide hydratase activity (Li et al., 2015).

MMS- and CY-induced *DDI2/3* expression is mediated by a zinc-finger transcription factor Fzf1 (Lin et al., 2023), which serves as a key regulator of several chemically responsive genes, including *DDI2/3*, *SSU1*, *YHB1* and *YNR064C* (Sarver and DeRisi, 2005). Fzf1 was initially reported as a regulator of *SSU1*, which encodes a sodium sulfite transporter (Avram and Bakalinsky, 1997), and its broader role in stress-responsive gene regulation was demonstrated in a genome-wide expression study (Sarver and DeRisi, 2005). Among Fzf1-regulated targets, *YHB1* encodes a nitric oxide dioxygenase that contributes to cellular defense against oxidative and nitrosative stress (Zhao et al., 1996); its expression is strongly induced by nitric oxide (NO) and related reactive nitrogen species (Horan et al., 2006). *YNR064C*, encoding a protein homologous to epoxide hydrolases (Elfstrom and Widersten, 2005), remains poorly characterized, and its expression profile under chemical stresses has yet to be fully elucidated.

The Fzf1 recognition consensus sequence CS2 was identified as a conserved 20-bp sequence located within promoter regions of all known Fzf1-regulated genes (Sarver and DeRisi, 2005). Fzf1 has been shown to bind specifically to the CS2 sequence, as demonstrated *in vitro* using both yeast cell extracts and recombinant Fzf1 protein (Lin et al., 2023; Ma et al., 2026). These findings collectively establish CS2 as a sequence-specific binding target of Fzf1.

Given that Fzf1 functions as a coordinated transcription factor, we investigated whether it uniformly activates all its target genes in response to various chemical stimuli by measuring expression of all known Fzf1-regulated genes under treatments with different inducers, including CY, MMS, sodium sulfite and NO. Interestingly, Fzf1-regulated gene expression varied significantly depending on the combination of target genes and chemical treatments. For example, CY induced *DDI2/3* expression up to 1,000-fold, while induced other Fzf1 targets by 20-40-fold. Similarly, NO treatment led to a 60-fold induction of *YNR064C*, whereas the remaining genes were induced less than 20-fold (Du et al., 2024). On the other hand, purified recombinant Fzf1 binds labeled CS2 probes from different promoters with comparable affinity (Ma et al., 2026). These observations collectively suggest that, despite sharing a common regulatory element, Fzf1-regulated genes respond differently to specific chemical stimuli, possibly due to interactions between other *cis*- and *trans*-acting regulatory elements.

To investigate these unknown *cis*- and *trans*-acting factors, we took two approaches. Firstly, given that *DDI2*/*3* exhibit exceptionally high induction levels in response to CY, we hypothesized that their promoters contain additional *cis*-acting elements specific for CY- and MMS-induction and performed extensive promoter deletion analyses. Secondly, a genome-wide microarray study revealed that under a histone H4 depletion condition, *DDI2* and *DDI3* were among the most highly induced genes (up to 300-fold), whereas *YNR064C* was induced by 20-fold and *YHB1* was repressed by fivefold (Wyrick et al., 1999) (Table S1). Based on the above observations, we hypothesized that nucleosome structures at these gene loci contribute to the differential expression of Fzf1-regulated genes in response to chemical stresses. To test this hypothesis, we utilized a previously reported system (Mann and Grunstein, 1992) to examine basal and chemical-induced expression of Fzf1-regulated genes under histone H3/H4-depleted conditions. These studies collectively revealed that *DDI2/3* are strongly repressed by their promoter nucleosome structures, and that CY and MMS derepress *DDI2/3* expression partially through histone depletion.

## Results

### Mapping *cis*-acting regulatory elements in the *DDI2* promoter

We have previously used plasmid YEp*_DDI2_*-*lacZ* to define CS2 sequence as a UAS, as an internal 27-bp deletion of CS2 abolished CY-and MMS-induced *DDI2-lacZ* expression (Lin et al., 2023). In this study, we made a series of internal and 5’ deletions in the *DDI2* promoter and measured their basal and chemical-induced transcript levels. The 5’ deletions revealed gradual increase in basal-level expression up to -229 (Figure 1A, B), which was accompanied by corresponding decrease in the fold induction (Figure 1A, C). Further deletion of the CS2 sequence (nt. -229 to - 190) did not alter the high basal transcript level, indicating that the chemical induction of *DDI2/3* expression mediated by the Fzf1-CS2 interaction at the *DDI2/3* promoter is exclusively achieved by de-repression. This high basal-level expression was maintained in subsequent deletions until hitting the TATA-box located at nt. -83 to -88 (5’-TATAAA-3’, Figures 1A and S1). Hence, the TATA box alone is required and appears to be sufficient to support the high-level *DDI2/3* expression.

**Figure 1.**
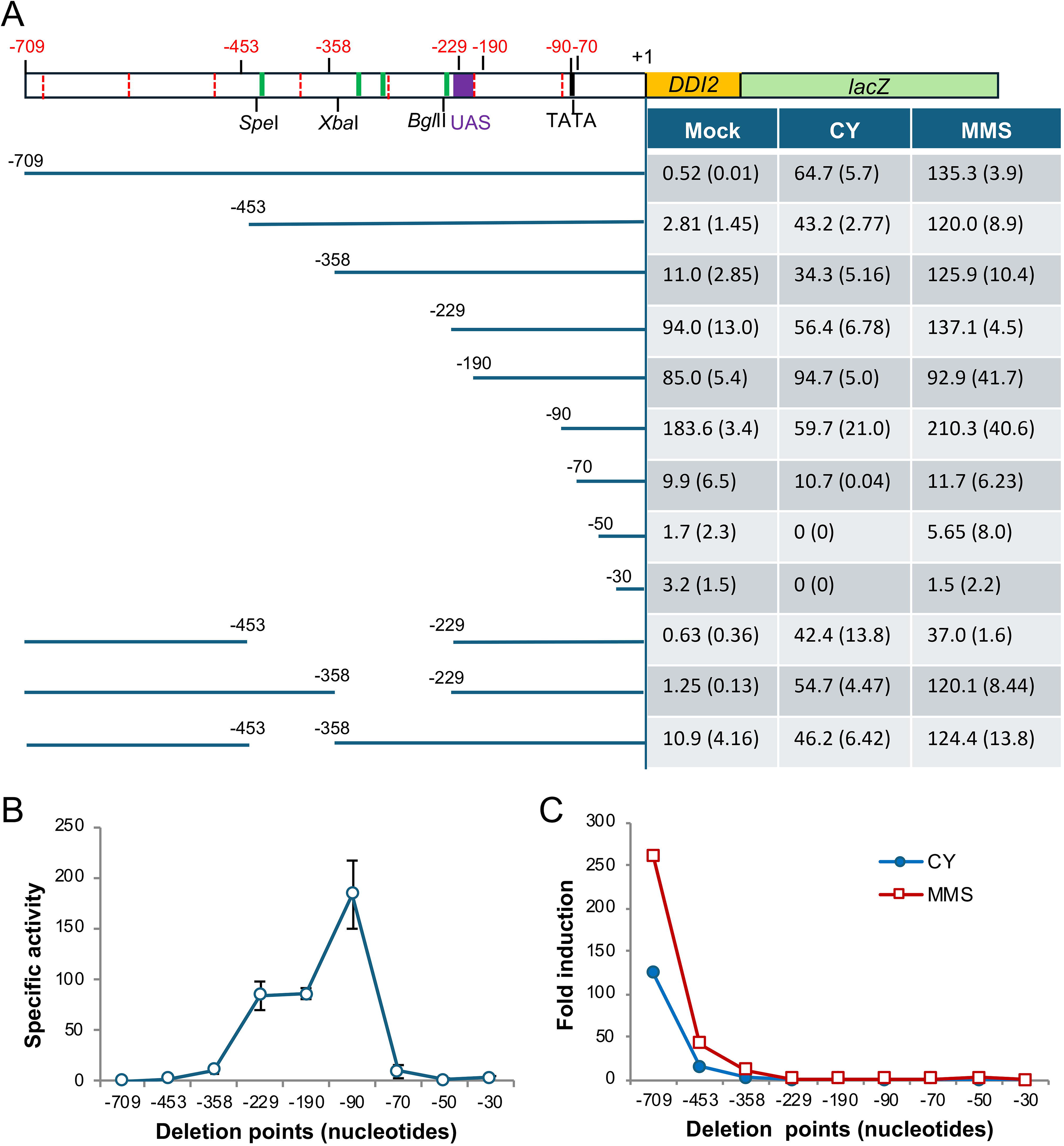
Identification of *cis*-acting elements in the *DDI2* promoter. **(A)** A series of promoter truncations were generated in plasmid YEp*_DDI2_*-*lacZ* and transformed into BY4741 cells. The left panel illustrates DNA sequence remaining after the truncation with marked truncation sites. The right panel shows the β-Gal activity of BY4741 cells carrying the indicated *P_DDI2_*-*lacZ* derivatives with CY, MMS or mock treatment. (B) Graphic illustration of basal level β-Gal activity of BY4741 cells carrying the indicated P*_DDI2_*-*lacZ* 5’ deletions. (C) Graphic illustration of fold induction of BY4741 cells carrying the indicated P*_DDI2_*-*lacZ* 5’ deletions in response to 20 mM CY (blue) or 0.1% MMS (red) treatment for 2 hours. Results of all β-Gal assays were from at least three independent experiments with standard deviations as shown in brackets.

The 5’ and internal deletions in the *DDI2* promoter indicate multiple URS elements within nt. -709 to -229 that cooperatively repress *DDI2/3*. Interestingly, the *DDI2* promoter sequence analysis revealed a 6-bp sequence, 5’-AAAAGA-3’, repeated four times within this region, located at positions -572 to -567, -331 to -326, -308 to -303 and -233 to -228 (Figures 1A and S1). Since a 6-nt sequence in DNA is typically found once every 4 kb, such four repeats within a 350-bp region indicate their significance and could explain the observed gradual derepressing effects. To test this possibility, we made sequential 6-bp deletions of the 5’-AAAAGA-3’ repeats and measured basal and CY-induced *DDI2-lacZ* transcript levels in the transformed cells. Figure S2 showed that all single and combined deletions did not affect CY-induced *DDI2-lacZ* expression, which effectively rules out their roles as URS elements.

An alternative approach was employed by searching for the Saccharomyces Genome Database (https://www.yeastgenome.org) to predict candidate transcription repressor binding sequences in the *DDI2* promoter. This analysis identified 15 putative binding sites for 15 transcription repressors within nt. -709 to -229 (Figure S3A). Among them, two transcription factors, Hsf1 and Ste12, were excluded from further investigation because their corresponding genes are essential to maintain cell viability, rendering their deletion lethal. The remaining 13 gene deletion strains, *ace2*, *swi5*, *adr1*, *ash1*, *bas1*, *gcn4*, *crz1*, *hac1*, *mot3*, *nrg1*, *rgt1*, *stb5* and *tec1*, were collected and transformed with the YEp*_DDI2_-lacZ* reporter plasmid. The expression *P_DDI2_*-*lacZ* gene was then measured following treatment with 20 mM CY for 2 hours. As shown in Fig. S3B, deletion of any of the 13 candidate transcription factor genes did not result in a dramatic increase in either basal-level or CY-induced reporter gene expression, suggesting that these genes do not play a major role in the *DDI2/3* regulation. However, one cannot rule out a possibility that these transcription factors are involved in the *DDI2/3* regulation, but function redundantly or indirectly.

### CY and MMS treatments cause histone depletion

Two pieces of information drew our attention to the nucleosome structure at *DDI2/3* promoters as an underlying mechanism of *DDI2/3* repression. Firstly, a study of chromosomal landscape on nucleosome-dependent gene expressions (Wyrick et al., 1999) revealed that *DDI2* and *DDI3* were among the most highly upregulated genes upon histone H4 depletion, thereby implicating a potential role of nucleosome occupancy in the *DDI2/3* expression. Secondly, search the *Saccharomyces NucMap* database (http://bigd.big.ac.cn/nucmap) (Oberbeckmann et al., 2019; Zhao et al., 2019) for the nucleosome distribution revealed that *DDI2/3* nt. -700 to - 250 regions are occupied by three nucleosomes (Figure 2A), which substantially overlap with our mapped URS elements (Figure 1).

**Figure 2.**
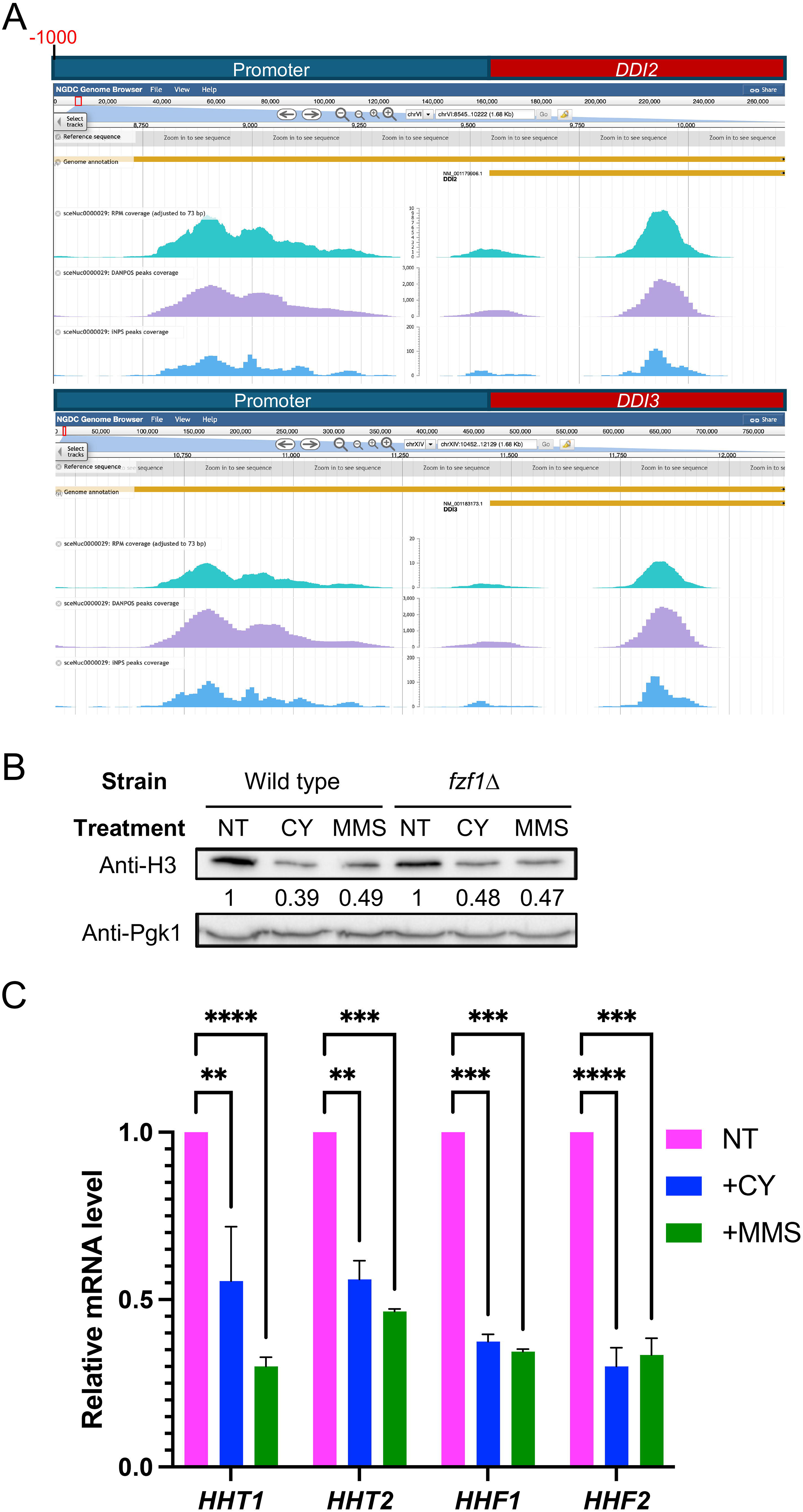
Effects of CY and MMS treatments on cellular histone levels. (A) Absolute nucleosome occupancy maps at the *DDI2* (upper panel) and *DDI3* (lower panel) promoter (up to 1 kb) and coding regions. Data were extracted from the *Saccharomyces NucMap* database (http://bigd.big.ac.cn/nucmap). RPM coverage provides a straightforward and rapid means to visualize overall nucleosome occupancy, DANPOS offers a robust framework for identifying and comparing dynamic chromatin features, and iNPS enables high-resolution nucleosome positioning. These three nucleosome mapping methods are included because *NucMap* database contains sufficient data for these approaches across a range of strains and experimental treatments. All datasets shown here are derived from strain BY4741, serving as the reference for comparisons with deletion strains. (B) Western blot analysis of histone H3 protein levels in BY4741 and its *fzf1*Δ mutant cells after 2-hour treatment with 20 mM CY or 0.1% MMS. H3 was detected using an anti-H3 antibody, with Pgk1 as an internal control. Numbers underneath the anti-H3 blot indicate relative histone H3 levels. (C) RT-qPCR analysis of genes encoding histone H3 (*HHT1* and *HHT2*) and H4 (*HHF1* and *HHF2*) after 2-hour treatment of BY4741 cells with 20 mM CY or 0.1% MMS. *UBC6* was used as the internal reference gene. Values represent the average of at least three biological replicates, with standard deviations shown as error bars. The statistical analysis was performed using two-way ANOVA multiple comparison via GraphPad Prism. ****, *P*<0.0001; ***, *P*< 0.001; **, *P*<0.01. NT, no chemical treatment.

Since CY and MMS treatments strongly and specifically induce *DDI2/3* expression (Du et al., 2024; Li et al., 2015; Lin et al., 2018), we hypothesized that CY and MMS induce *DDI2/3* expression through histone depletion. To test this hypothesis, we treated BY4741 cells with 20 mM CY or 0.1% MMS for 2 hours, which were conditions previously optimized for the *DDI2/3* induction, followed by western blot analysis against an anti-histone H3 antibody. Figure 2B shows that CY and MMS treatments reduced cellular histone H3 protein levels by more than 50%. Furthermore, deletion of *FZF1* did not impact on CY/MMS-induced histone H3 depletion (Figure 2B), indicating that this effect is independent of Fzf1-mediated transcriptional regulation. To ask whether these inhibitory effects occur at the transcriptional, translational or post-translational level, we measured mRNA levels of both copies of histone *H3* and (*HHT1* and *HHT2*) and *H4* (*HHF1* and *HHF2*) genes by quantitative RT-PCR (RT-qPCR) analysis, which revealed that CY and MMS treatments reduced both *H3* and *H4* transcript levels by 50% or more, and the differences were statistically significant (Figure 2C). Together, these findings suggest that CY and MMS treatments enhance *DDI2/3* expression by decreasing cellular histone abundance and hence modulating chromatin structures at *DDI2/3* promoters.

### Effects of *spt10*-mediated histone depletion on the *DDI2/3* expression

If the histone depletion caused by CY or MMS alone is sufficient to induce the massive *DDI2/3* expression, other means of histone depletion can also induce *DDI2*/*3* expression. Spt10 functions as a transcription activator that specifically recognizes tandem UAS elements within core histone gene promoters, thereby promoting histone gene expression (Eriksson et al., 2012). We reasoned that deletion of *SPT10* could reduce histone gene expression, leading to histone depletion reminiscent of CY and MMS treatment, which allowed us to address roles of histone depletion in the CY/MMS-induced *DDI2/3* expression. To this end, we treated BY4741 *spt10*Δ cells with or without 20 mM CY for 2 hours and measured histone *H3* and *H4* transcript levels by RT-qPCR. Figure 3A showed that deletion of *SPT10* reduced all four histone gene transcripts to levels comparable to the CY treatment, while CY treatment of *spt10*Δ cells did not affect *HHT1* expression but further reduced *HHT2*, *HHF1* and *HHF2* transcript levels. As anticipated, cellular histone H3 protein levels were markedly reduced in *SPT10*Δ cells, while CY or MMS treatment did not further deplete histone H3 in *spt10*Δ cells.

**Figure 3.**
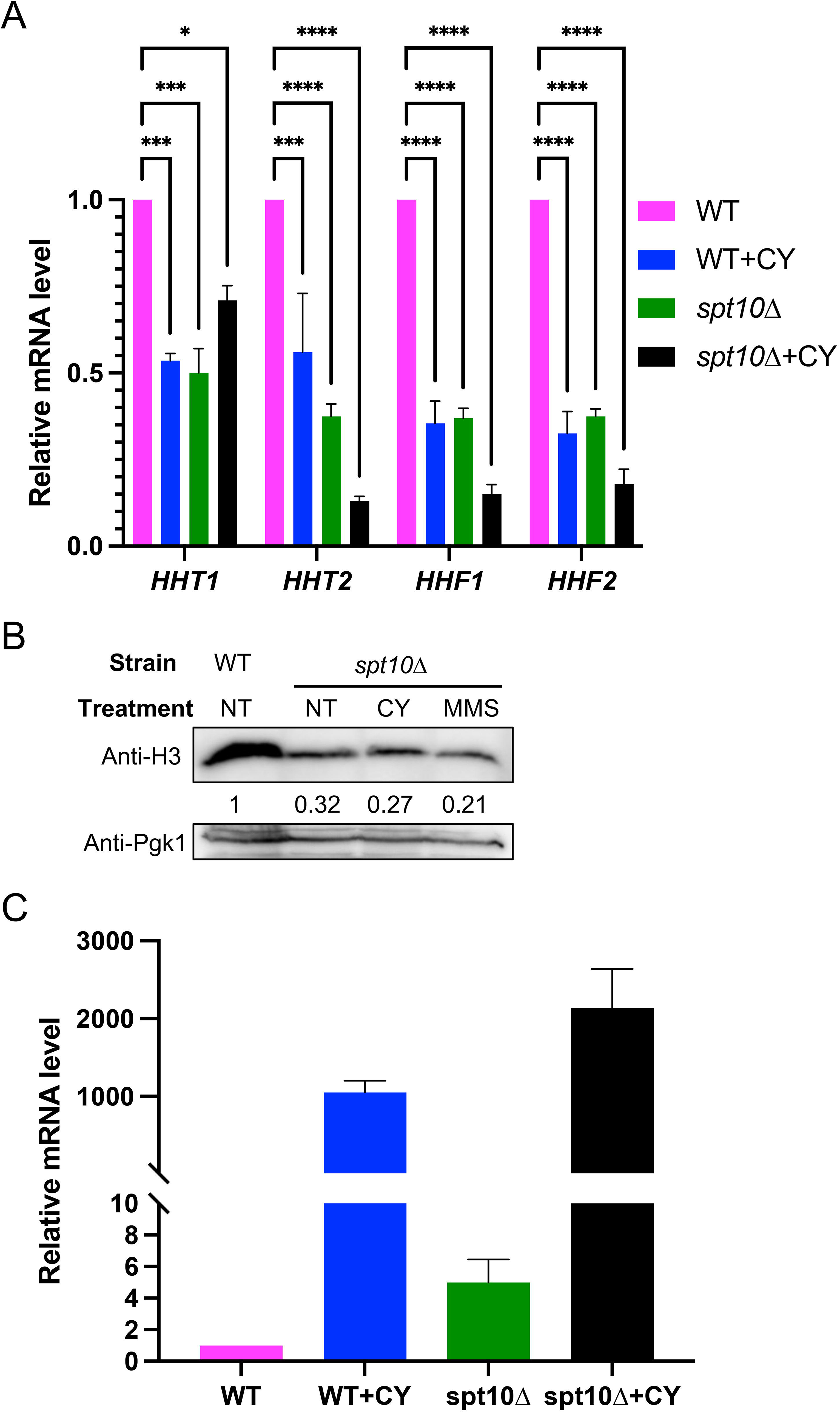
Effects of *SPT10* deletion on histone and *DDI2/3* gene expression. (A) RT-qPCR analysis of genes encoding histone H3 (*HHT1* and *HHT2*) and H4 (*HHF1* and *HHF2*) after 2-hour treatment of BY4741 (WT) or BY4741 *spt10*Δ cells with 20 mM CY. *UBC6* was used as the internal reference gene. (B) Western blot analysis of histone H3 protein levels in BY4741 and its *spt10*Δ mutant cells after 2-hour treatment with 20 mM CY or 0.1% MMS. H3 was detected using an anti-H3 antibody, with Pgk1 as an internal control. Numbers underneath the anti-H3 blot indicate relative histone H3 levels. (C) RT-qPCR analysis of *DDI2/3* transcript levels in *spt10*Δ cells with or without chemical treatment. The *DDI2/3* transcript were measured in the same sets of samples as in (A). Values in both (A) and (C) represent the average of at least three biological replicates, with standard deviations shown as error bars. The statistical analysis was performed using two-way ANOVA multiple comparison via GraphPad Prism. ****, *P*<0.0001; ***, *P*< 0.001; *, *P*<0.05. NT, no chemical treatment.

Subsequently, we examined *DDI2/3* expression in *spt10*Δ cells with and without CY treatment and compared it to wild-type cells. As shown in Figure 3C, deletion of *SPT10* alone caused a fivefold increase in the *DDI2/3* expression. In comparison, CY treatment of wild-type cells induced *DDI2/3* expression by approximately 1,000-fold, while in *spt10*Δ cells, CY treatment induced *DDI2/3* by approximately 2,000-fold. These findings collectively suggest that histone depletion alone, although capable of moderately inducing *DDI2/3*, cannot account for the observed massive *DDI2/3* induction by CY and MMS.

### CY and MMS affect yeast cell growth via different mechanisms

MMS is a well-known methylating agent that methylates macromolecules including DNA, RNA and proteins (Zhang et al., 2005). Its roles as a DNA-damaging agent have been extensively characterized (Beranek, 1990). As a genotoxic agent, MMS causes DNA adducts that block replication, leading to cell death (Wyatt and Pittman, 2006). However, CY exhibits very mild if any toxicity in yeast cells (Lin et al., 2018), and its mechanistic effects remain unclear. To systematically assess how yeast cells respond to these chemical stresses, we monitored BY4741 cell growth in the presence of different concentrations of CY and MMS for two hours followed by measuring cell density at OD□□□ _nm_. Figure 4A shows that within the experimental concentration ranges, CY and MMS inhibited cell proliferation at comparable rates. For example, 40 mM CY reduced cell density by 75%, while 0.1% MMS reduced cell density by 70%.

**Figure 4.**
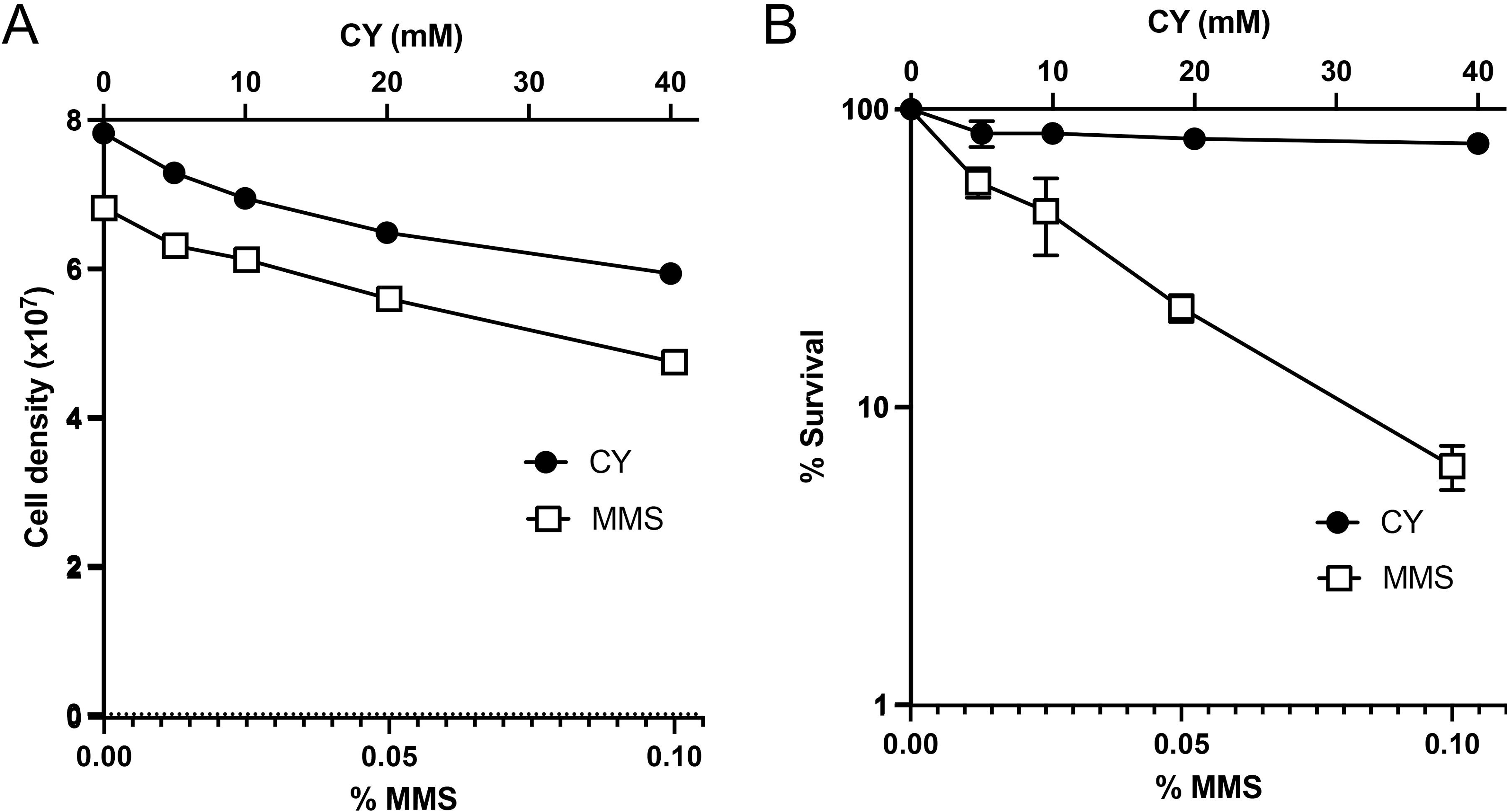
Effects of CY and MMS treatments on yeast cell growth and survival. (A) Cell growth as measured by cell density. (B) Cell survival as measured by % survival. BY4741 cells were cultured to mid-log phase (OD _nm_ = 0.4 - 0.5) and subsequently treated with 5, 10, 20 or 40 mM CY, or with 0.0125%, 0.025%, 0.05% or 0.1% MMS for 2 hours. After treatment, cell density was measured by spectrophotometry at OD _nm_, and aliquots of the cultures were serially diluted and plated on YPD agar plates. Survival rates were calculated by comparing the number of colony-forming units (CFUs) from treated samples to those from untreated controls. Values in both (A) and (B) represent the average of at least three biological replicates, with standard deviations shown as error bars.

To evaluate the impact of CY and MMS treatments on cell viability, we treated BY4741 cells at indicated chemical concentrations for two hours followed by dilution and plating cells on non-selective YPD plates. After incubation for two days at 30 °C, number of colonies on each plate were counted and cell viability was calculated based on percentage colony-forming units. MMS and CY treatments resulted in dramatic difference in cell viability. As shown in Figure 4B, after 40 mM CY treatment for 2 hours, 77% cells were viable and formed colonies. In contrast, after 0.1% MMS treatment for two hours, only 6% cells were viable and formed colonies. These observations support a notion that, unlike MMS, which is highly toxic to yeast cells, CY primarily inhibits cell division without significantly compromising overall cell viability under our experimental conditions.

### Histone depletion differentially affects Fzf1-regulated gene expression

To critically test a hypothesis that nucleosome positioning in the *DDI2/3* promoters serves as *cis*-acting URS, we wished to recapitulate the histone H4 depletion conditions and measure *DDI2/3* expression. Unfortunately, the reported yeast strain became unavailable from original authors. Instead, we used a strain RMY102 (Mann and Grunstein, 1992), which was constructed by introducing a plasmid carrying *HHT2* driven by a *GAL10* promoter (*P_GAL10_*-*HHT2*) and *HHF2* driven by a *GAL1* promoter (*P_GAL1_*-*HHF2*) and then deleting the endogenous *HHT1/2* and *HHF1/2* genes encoding histone H3 and H4, respectively. When growing a YPGal medium in which galactose is the sole carbon source, RMY102 cells express both *HHT2* and *HHF2* genes and exhibit characteristic growth. Upon switching to a YPD glucose-containing medium, *P_GAL10_*-*HHT2* and *P_GAL1_*-*HHF2* genes are repressed, leading to a histone depletion state and gradual decline in cell growth. This experimental setup is illustrated in Figure 5A.

**Figure 5.**
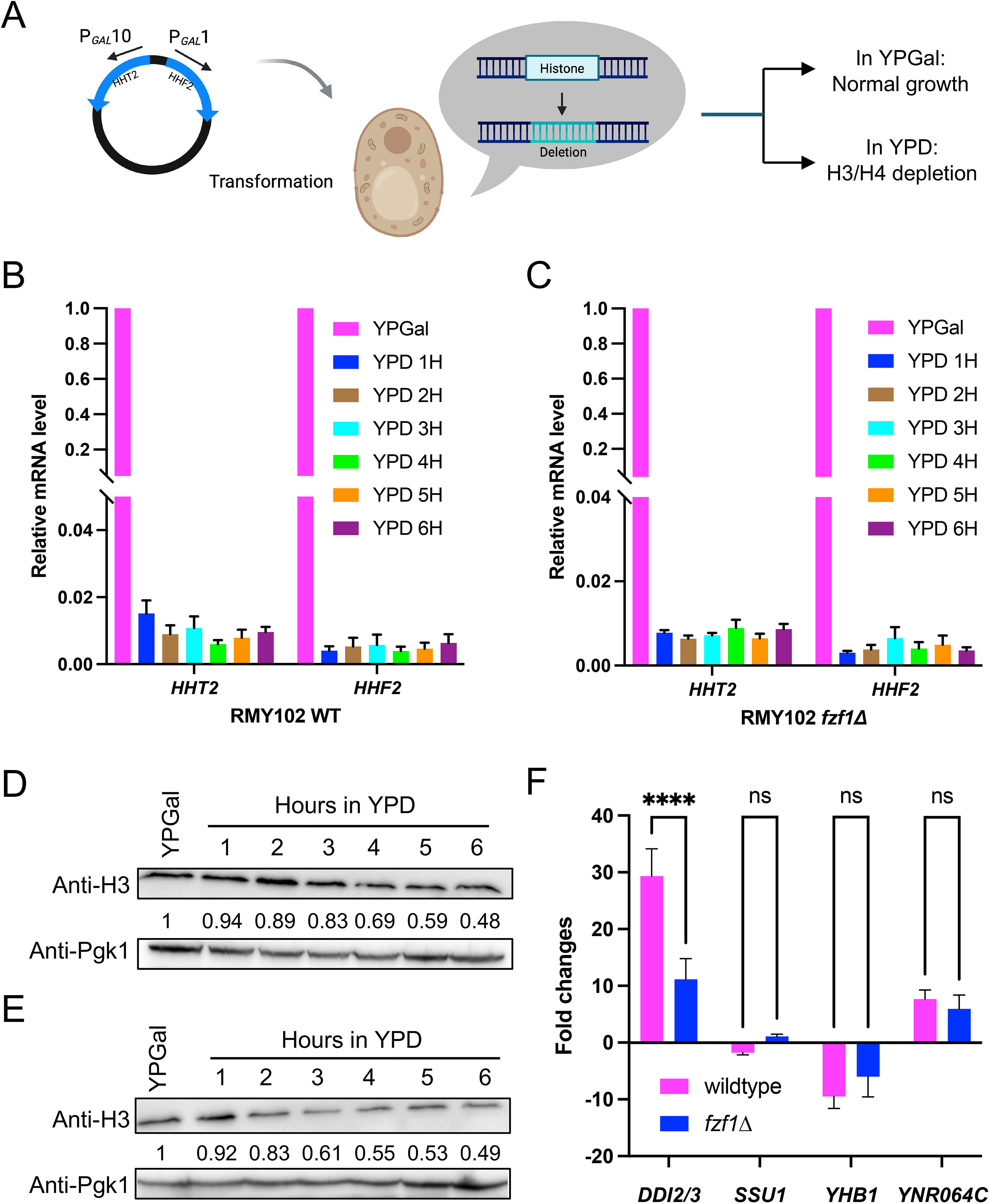
Histone depletion and its effects on the expression of Fzf1-regulated genes. (A) Schematic diagram illustrating the histone depletion strain RMY102 and the experimental plan. Both copies of genes encoding histone H3 and H4 are deleted, and RMY102 cells are maintained by a plasmid carrying *HHT2* and *HHF2* under the control of galactose-inducible promoters *P_GAL_10* and *P_GAL_1*, respectively, and incubation in YPGal medium. Once transferring to a YPD medium, the *P_GAL_* promoters are repressed, leading to cessation of *de novo* H3 and H4 synthesis and creation of a histone depletion status. (B,C) RT-qPCR analysis of cellular *HHT2* and *HHF2* levels in RMY102 (B) and RMY102 *fzf1*Δ (C) cells under H3/H4 depletion conditions. (D, E) Western blot analysis of cellular histone H3 levels in histone depleted RMY102 (D) and RMY102 *fzf1*Δ (E) cells over time. Whole-cell lysates from yeast cells grown in either YPGal or YPD medium for the indicated times were subjected to immunoblotting using anti-H3 antibodies (upper panel), and anti-Pgk1 (lower panel) served as an internal control. Numbers underneath the anti-H3 blot indicate histone H3 levels relative to cells grown in YPGal. (F) RT-qPCR analysis of transcript levels of Fzf1-regulated genes in RMY102 and RMY102 *fzf1*Δ under histone depletion conditions. Values represent the average of at least three biological replicates, with standard deviations shown as error bars. The statistical analysis was performed using two-way ANOVA multiple comparison via GraphPad Prism. ****, *P*<0.0001; ns, not statistically significant.

To monitor the histone depletion dynamics in RMY102 cells, we measured *HHT2* and *HHF2* transcript levels in RMY102 cells by RT-qPCR. As shown in Figure 5B, upon shifting from YPGal to the YPD medium, both transcript levels dropped rapidly within first hour and remained at or less than 1% in comparison to cells in the YPGal medium. Similar *HHT2* and *HHF2* mRNA depleting effects were also observed in *FZF1*-deleted RMY102 cells (Figure 5C). Consequently, H3 protein levels were reduced in RMY102 (Figure 5D) and RMY102 *fzf1*Δ (Fig. 5E) cells in YPD compared to corresponding YPGal-grown cells. After incubating 6 hours in YPD, cell density increased by twofold (Figure S4), which was accompanied with 50% decrease in cellular H3 levels (Figure 5D, E), indicating that *de novo* histone H3 and H4 synthesis was stopped shortly after medium shift, and cellular H3 and H4 levels were diluted primarily through cell division, marking cells in a well-defined histone depletion state. We then assessed the expression of Fzf1-regulated genes in histone-depleted RMY102 cells. As shown in Figure 5F and Table S2, after 6-hour incubation in YPD, *DDI2/3* were induced to nearly 30-fold, and *YNR064C* was also induced by nearly eightfold. Meanwhile, the *SSU1* transcript level remained unchanged, while *YHB1* was repressed by tenfold. To ask whether these changes were dependent on Fzf1, we repeated the histone depletion experiment in RMY102 *fzf1*Δ cells. As shown in Figure 5F, *DDI2/3* and *YNR064C* were still induced upon histone depletion, although the *DDI2/3* induction in *fzf1*Δ was reduced to tenfold, while deletion of *FZF1* did not further affect *SSU1*, *YHB1* and *YNR064C* expression. These results, together with a previous genome-wide microarray data (Wyrick et al., 1999), collectively indicate that histone depletion differentially affects Fzf1-regulated genes; it induces *DDI2/3* and *YNR064C*, represses *YHB1*, and has little effect on the *SSU1* expression. Furthermore, histone depletion affects Fzf1-regulated genes largely in an Fzf1-independent manner.

Under the above histone depletion conditions, CY and MMS treatments further reduced cellular H3 levels (Figure S5), indicating joint effects of experimental histone depletion and CY/MMS treatment on the Fzf1-regulated gene expression.

### Effects of H3/H4 depletion on the expression of Fzf1-regulated genes in response to CY

We further investigated effects of the CY treatment on Fzf1-regulated gene expression under histone depletion conditions. RMY102 cells were cultured in a YPD medium to deplete histones H3 and H4 for 4 hours, followed by an additional 2-hour CY treatment. As shown in Figure 6A, in comparison to untreated RMY102 cells grown in YPGal, as low as 5 mM CY induced *DDI2/3* expression by 350-fold under histone depletion conditions and further increase in CY concentrations did not result in corresponding induction of *DDI2/3*. In comparison, in BY4741 cells, the *DDI2/3* expression was induced by 5 mM CY to 285-fold and by 20 mM CY to 1,050-fold in wildtype cells without histone depletion, and other Fzf1-regulated genes were also induced by CY in a dose-dependent manner in BY4741 cells (Du et al., 2024); however, under the histone depletion conditions, their responses were barely dose-dependent (Figure 6A).

**Figure 6.**
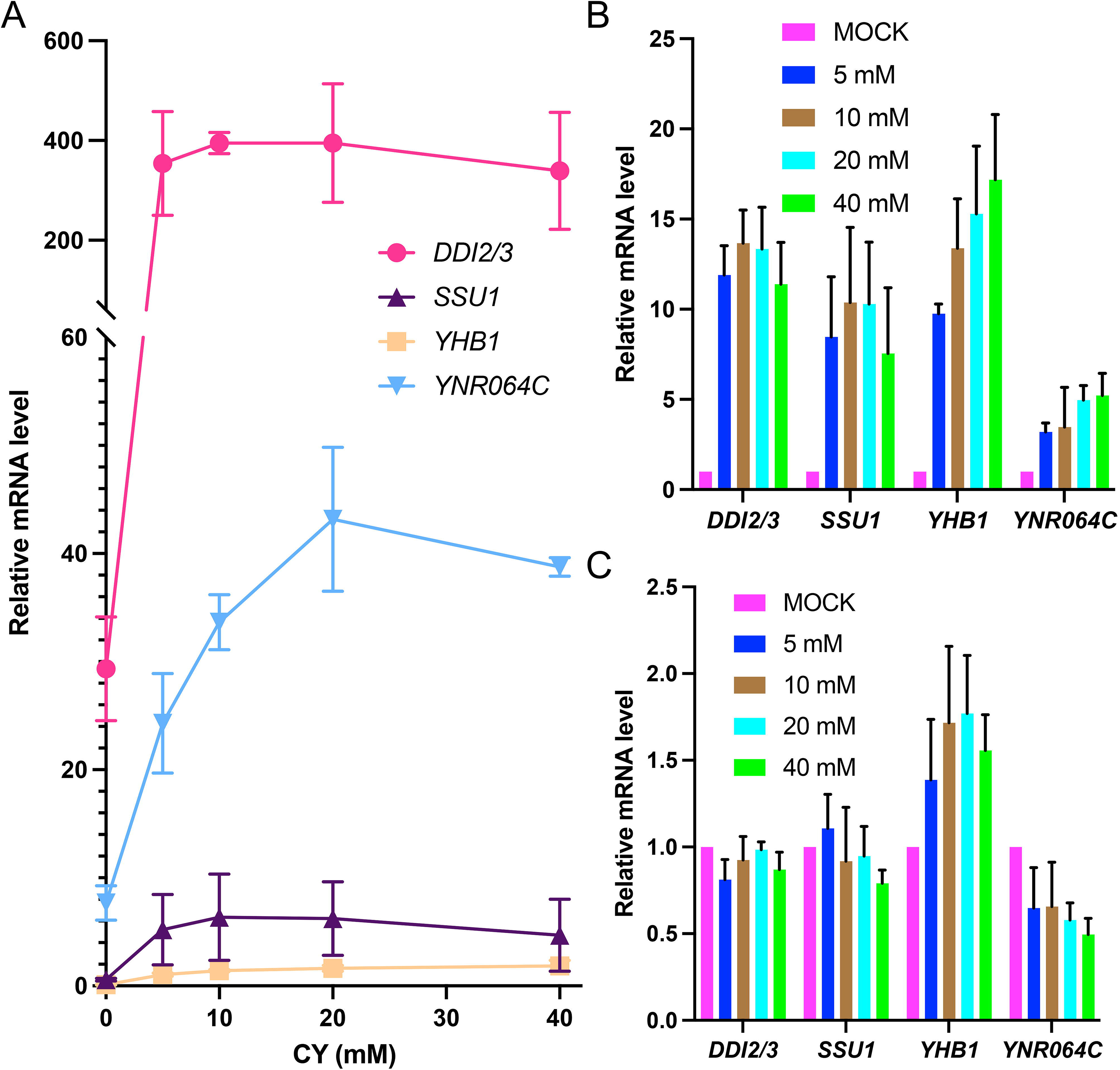
Joint effects of CY treatment and histone depletion on the expression of Fzf1-regulated genes. (A) Relative mRNA levels of Fzf1-regulated genes in RMY102 cells treated with different concentrations of CY under histone depletion conditions. Cells without histone H3/4 depletion and CY treatment was set up as “1” (not showing in the figure). Gene expression was quantified by RT-qPCR. Data represent the mean values from at least three independent biological replicates, with standard deviations represented as error bars. (B) Comparative analysis of Fzf1-regulated gene expression in RMY102 cells under histone H3/4 depletion with various CY treatment conditions. Cells under the histone H3/4 depletion condition without CY treatment was set up as “1” (magenta bars). (C) Relative mRNA levels of Fzf1-regulated genes in RMY102 *fzf1*Δ cells under histone H3/H4 depletion with various CY treatment conditions. Cells under the histone H3/4 depletion condition without CY treatment was set up as “1” (magenta bars).

The Fzf1-regulated genes are known to be variably indued by CY treatments in BY4741 cells, in which *DDI2/3* are induced by 1,000-fold and *YHB1* by less than 20-fold (Du et al., 2024). This study revealed that these genes were also differentially affected by nucleosome structures, as evident by their expression under histone depletion conditions. Interestingly, when the histone depletion effects are taken out as variables, *DDI2/3*, *SSU1* and *YHB*1 were induced by CY to comparable levels of approximately 10-15-fold, regardless of CY concentrations (Figure 6B). Similarly, *YNR064C* was induced by CY up to fivefold in a dose-dependent manner (Figure 6B), indicating that this gene has its own unique regulatory components.

We also assessed whether Fzf1 is still required for CY-mediated induction under histone depletion conditions. As shown in Fig. 6C, deletion of *FZF1* completely abolished CY-induced expression of all tested genes under histone-depleted conditions, suggesting that Fzf1 plays an essential role in CY-induced transcriptional activation independently of its role(s) in combating nucleosome effects.

### Effects of H3/H4 depletion on the expression of Fzf1-regulated genes in response to MMS

Similarly, after incubating RMY102 cells in a YPD medium for 4 hours, we treated cells with varying concentrations of MMS for 2 hours and then measured transcript levels of Fzf1-regulated genes. As shown in Figure 7A, under the above histone depletion conditions, MMS treatments led to variable levels of Fzf1-regulated gene expression compared to untreated and non-depleted RMY102 cells grown in YPGal. Like CY treatment, increasing MMS concentrations did not further induce these genes and even led to a moderate reduction in the *DDI2/3* expression, which contrasted with previously observed dose responses in BY4741 cells (Du et al., 2024). To separate the effect of MMS treatments from the nucleosome impact, we again compared transcript levels of histone H3/H4 depleted RMY102 cells with and without MMS treatment. As shown in Figure 7B, all Fzf1-regulated genes were induced by MMS to comparable levels with no observable dose responses. We also assessed the expression of Fzf1-regulated genes in RMY102 *fzf1*Δ cells under histone depletion and MMS treatment conditions. As shown in Figure 7C, deletion of *FZF1* abolished MMS-induced gene expression, indicating that *FZF1* is required for MMS-mediated transcriptional activation under histone-depleted conditions.

**Figure 7.**
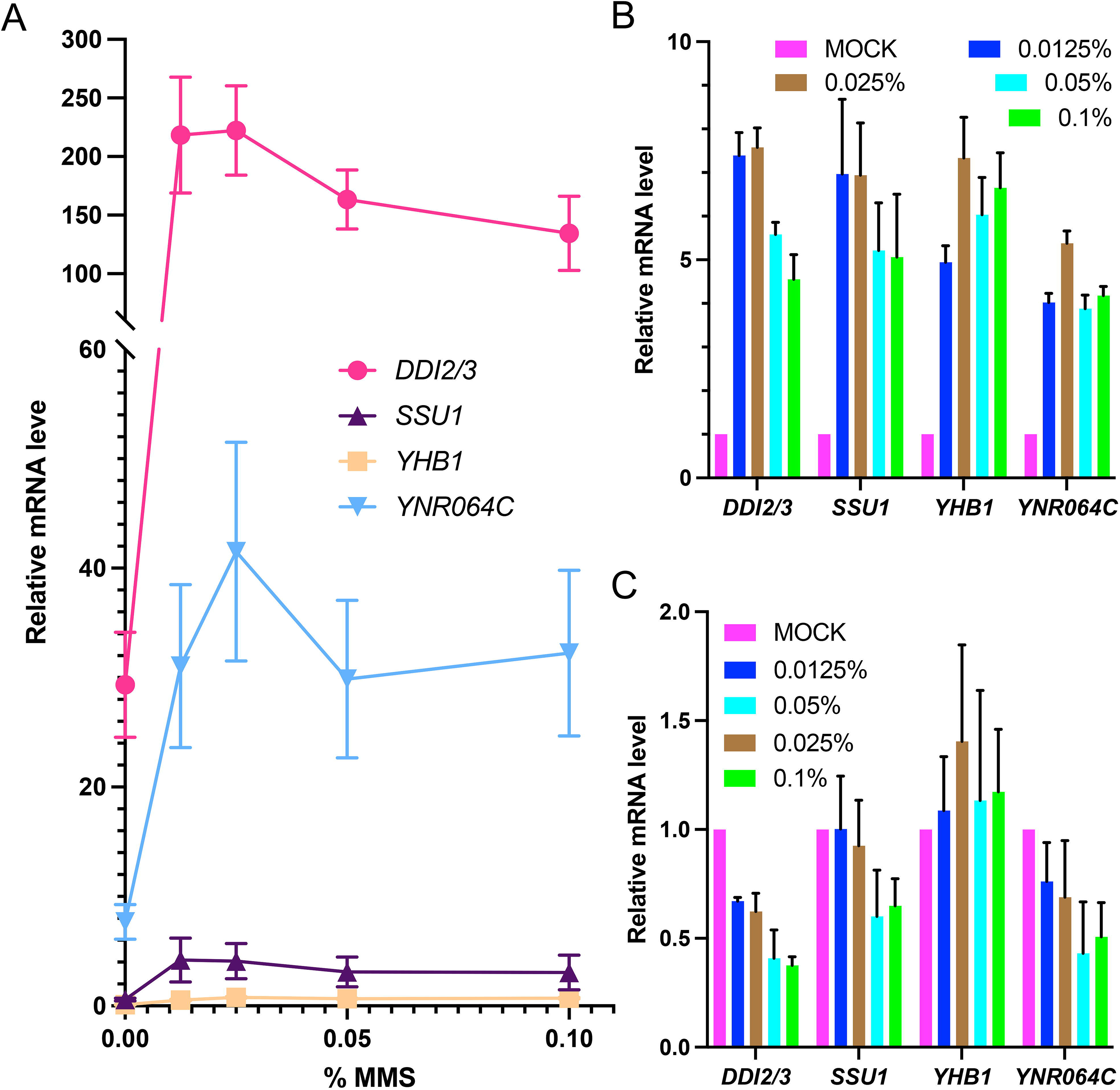
Joint effects of MMS treatment and histone depletion on the expression of Fzf1-regulated genes. (A) Relative mRNA levels of Fzf1-regulated genes in RMY102 cells treated with different concentrations of MMS under histone depletion conditions. Cells without histone H3/4 depletion and MMS treatment was set up as “1” (not showing in the figure). Gene expression was quantified by RT-qPCR. Data represent the mean values from at least three independent biological replicates, with standard deviations represented as error bars. (B) Comparative analysis of Fzf1-regulated gene expression in RMY102 cells under histone H3/4 depletion with various MMS treatment conditions. Cells under histone H3/4 depletion condition without MMS treatment was set up as “1” (magenta bars). (C) Relative mRNA levels of Fzf1-regulated genes in RMY102 *fzf1*Δ cells under histone H3/H4 depletion with various MMS treatment conditions. Cells under histone H3/4 depletion condition without MMS treatment was set up as “1” (magenta bars).

### Effects of CY on nucleosome occupancy in *DDI2/3* promoters

To directly assess whether chemical treatments, particularly CY, affects nucleosome occupancy at *DDI2/*3 promoter regions, we performed MNase-seq to interrogate chromatin structures *in vivo*. BY4741 and its *fzf1*Δ mutant cells were treated with or without 20 mM CY for 2 hours, after which nucleosome occupancy was profiled globally, and we focused on the *DDI2/3* coding and upstream sequences in this study. As the CY-induced depletion of H3/H4 levels was predicted to result in reduced genome-wide nucleosome occupancy, we normalized all data sets to genomic mean (RPGC) to identify those regions that showed increased nucleosome loss compared to the rest of the genome. MNase digestion profiles and the RPGC-normalized heatmaps are presented in Figure S6. Employing only sequence reads that uniquely mapped to either *DDI2* or *DDI3* showed that CY treatment of wild-type cells led to a marked decrease in the nucleosome occupancy upstream and within the coding sequences of both *DDI2* and *DDI3* (Figure 8A, B), which included three regions of high nucleosome density: nt. -1,000 to -200, - 200 to +160, and +250 to +550 relative to the translation start site. Importantly, the -1,000 to - 200 region overlaps with the previously defined *cis*-acting URS elements (Figure 1) and *NucMap* database analyses (Figure 2A), demonstrating that CY treatment *in vivo* can significantly reduce nucleosome occupancy across *DDI2/3* upstream regulatory regions. As expected, CY treatment in *fzf1*Δ cells failed to reduce nucleosome occupancy at *DDI2/3* promoter and coding sequences including the nt. -1,000 to -200 region (Figure 8A, B), suggesting that Fzf1 is required for the CY-mediated chromatin remodeling at these loci. Interestingly, the CY treatment also reduced nucleosome occupancy at promoter and coding regions of *SSU1* (Figure S7A) and *YHB1* (Figure S7B) in a Fzf1-dependent manner. In contrast, the CY treatment did not appear to affect the nucleosome occupancy at the *YNR064C* locus (Figure S7C), indicating that CY differently affects nucleosome structures of Fzf1-regulated genes. Based on the annotated *S. cerevisiae* nucleosome intervals (Weiner et al., 2015), we quantified nucleosome occupancy in the *DDI2* and *DDI3* promoters at the single nucleosome resolution in comparison to the mean of all other upstream nucleosomes, as shown in Figure 8C. While CY treatment and the *fzf1* mutation did not affect -1 nucleosome occupancy, CY treatment dramatically reduced nucleosome occupancy at - 2 and moderately reduced nucleosome occupancies at -3 and -4, all in an Fzf1-dependent manner (Figure 8C). These observations agree with the *DDI2* promoter deletion analysis (Figure 1), in which 5’ deletion gradually increased the basal level expression of *DDI2-lacZ* with the strongest effect observed by deletion of the -2 nucleosome binding sequence (nt -415 to -269).

**Figure 8.**
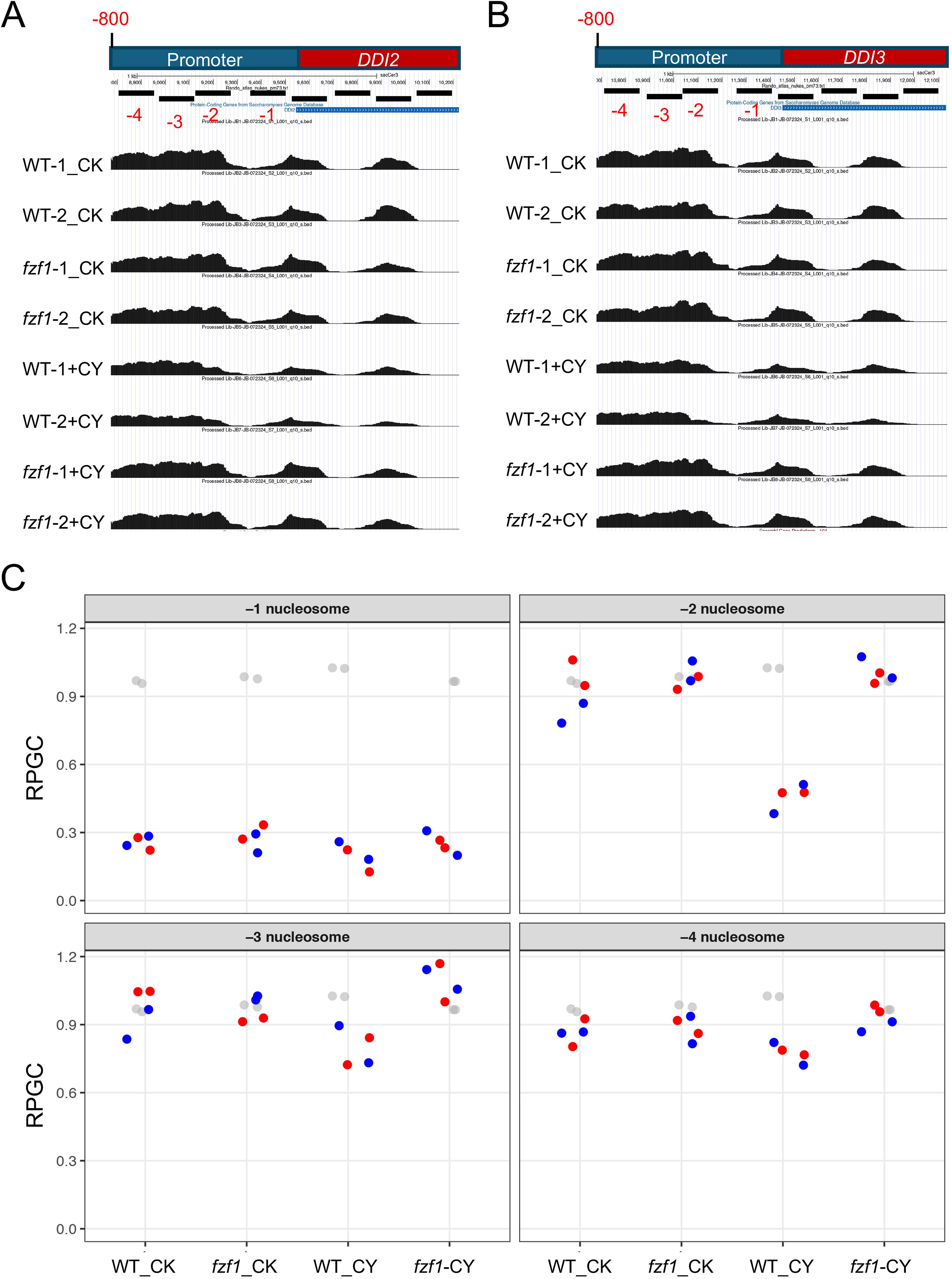
Nucleosome occupancy at the *DDI2* and *DDI3* loci under various experimental conditions. (A, B) Nucleosome occupancy profiles of (A) *DDI2* and (B) *DDI3* coding regions and up to 1 kp from the translation start in BY4741 (WT) and BY4741 *fzf1*Δ cells with or without 20 mM CY treatment for 2 hours. Two independent replicates were performed for each treatment. Only reads that uniquely mapped to *DDI2* or *DDI3* were used in the analysis, and the data were normalized to the genomic mean (Reads Per Genomic Content, RPGC). (C) The occupancy of individual nucleosomes upstream of *DDI2* and *DDI3* based on (Weiner et al., 2015). The four upstream nucleosomes in *DDI2* (blue) and *DDI3* (red) were compared with the mean of all other upstream (−1, -2, and -3) nucleosomes (grey) under given experimental conditions.

## Discussion

In this study, we investigated whether and how nucleosome structures play a role in the expression of Fzf1-regulated genes. Our central hypothesis was derived from *DDI2/3* promoter dissection and their expression in response to chemical stresses, and subsequent investigation helped to understand how other Fzf1-regulated genes are controlled by their nucleosome structures. This research led to several conclusions. Firstly, the *DDI2/3* expression is suppressed by promoter URS elements located at nt. -700 to -250, which is associated with heavy nucleosome occupancy, and their induction by CY/MMS is therefore mediated by Fzf1-dependent de-repression. The above conclusion agrees with our previous identification of Fzf1-ZF4 as a repressor element (Du et al., 2024), and further suggests that the Fzf1-ZF5 activation domain functions to counteract the nucleosome, as illustrated in Figure 9, instead of promoting the general transcription initiation as previously thought (Du et al., 2026). Secondly, the primary parameter contributing to different levels of chemical-induced responses among Fzf1-regulated genes is how they are controlled by their existing nucleosome structures under non-induced conditions. The nucleosome distribution patterns among Fzf1-regulated gene promoters are rather different, and consequences of histone depletion are also different. For example, histone depletion alone can induce *DDI2/3* by 30-fold (up to 300-fold in a previous report) while repress *YHB1* by tenfold, which could explain why CY can induce *DDI2/3* by 1,000-fold but *YHB1* by less than 20-fold. Thirdly, after the differential control of these gene expression by nucleosomes is taken into consideration, the remaining levels of chemical induction of these genes are comparable: around tenfold by CY and sixfold by MMS. This observation is consistent with our previous observations that purified recombinant Fzf1 protein binds different CS2 sequences with comparable affinities *in vitro* (Ma et al., 2026), indicating that the Fzf1-CS2 interaction in response to chemical stresses plays similar roles across these promoters. Fourthly, Fzf1 appears to be required for both de-repression through chromatin remodeling at *DDI2/3* promoters and activation in response to chemical stresses, as illustrated in Figure 9. These dually regulatory mechanisms have been dissected by two different means. One was revealed in studying *fzf1-ZF4* (Du et al., 2024) mutants, in which the basal-level Fzf1-regulated gene expression was elevated, while *DDI2/3* can still be induced by CY. However, deletion of *FZF1* completely abolishes *DDI2/3* induction (Du et al., 2024; Lin et al., 2023). Another was that, upon histone depletion, Fzf1-regulated genes could still be further induced by CY and MMS in an Fzf1-dependent manner. Fifthly, although both CY and MMS treatments can inhibit yeast cell growth and proliferation, and suppress histone gene expression, their modes of action differ to certain extend. MMS is a well-known methylating agent that can methylate macromolecules including proteins (Paik et al., 1984). Indeed, it has been previously reported that MMS can methylate the Fzf1-K70 residue *in vitro*, and that an *fzf1-K70A* mutation completely abolishes MMS-induced *DDI2/3* expression (Lin et al., 2023). It remains unknown whether histone subunits can be methylated, or their acetylation levels can be affected (Lee et al., 2007a), by MMS. CY can also modify a Lys residue to form homoarginine (Amberger, 2013), although such type of PTM and its biological significance has not yet been reported. It would be interesting to investigate whether the CY treatment induces PTMs in Fzf1 and/or histone subunits as an underlying mechanism of chemical signaling. On the other hand, a clear difference between CY and MMS treatments as revealed in this study is that MMS, as a well-known DNA-damaging agent, caused toxicity and cell death, while the CY treatment under our experimental conditions inhibited cell growth with little toxicity and cell death. The biological effects of CY deserve further investigation. Sixthly, although the nucleosome occupancy at both *DDI2/3* and *YHB1* promoters are reduced after the CY treatment, its biological consequences are different, since reducing cellular histone levels derepressed *DDI2/3* expression while inhibited *YHB1* expression. Similarly, other Fzf1-regulated genes may contain other unique *cis*-acting regulatory elements to govern their expression in response to chemical stresses. Finally, study of nucleosome occupancy at *DDI2/3* promoters at the single nucleosome resolution revealed that CY treatment dramatically reduced the nucleosome occupancy at -2 location, which correlates well with the *DDI2* promoter deletion analysis, indicating that the CY treatment releases this nucleosome in an Fzf1-dependent manner. How chromatin structures affect other Fzf1-regulated gene expression requires further investigation.

**Figure 9.**
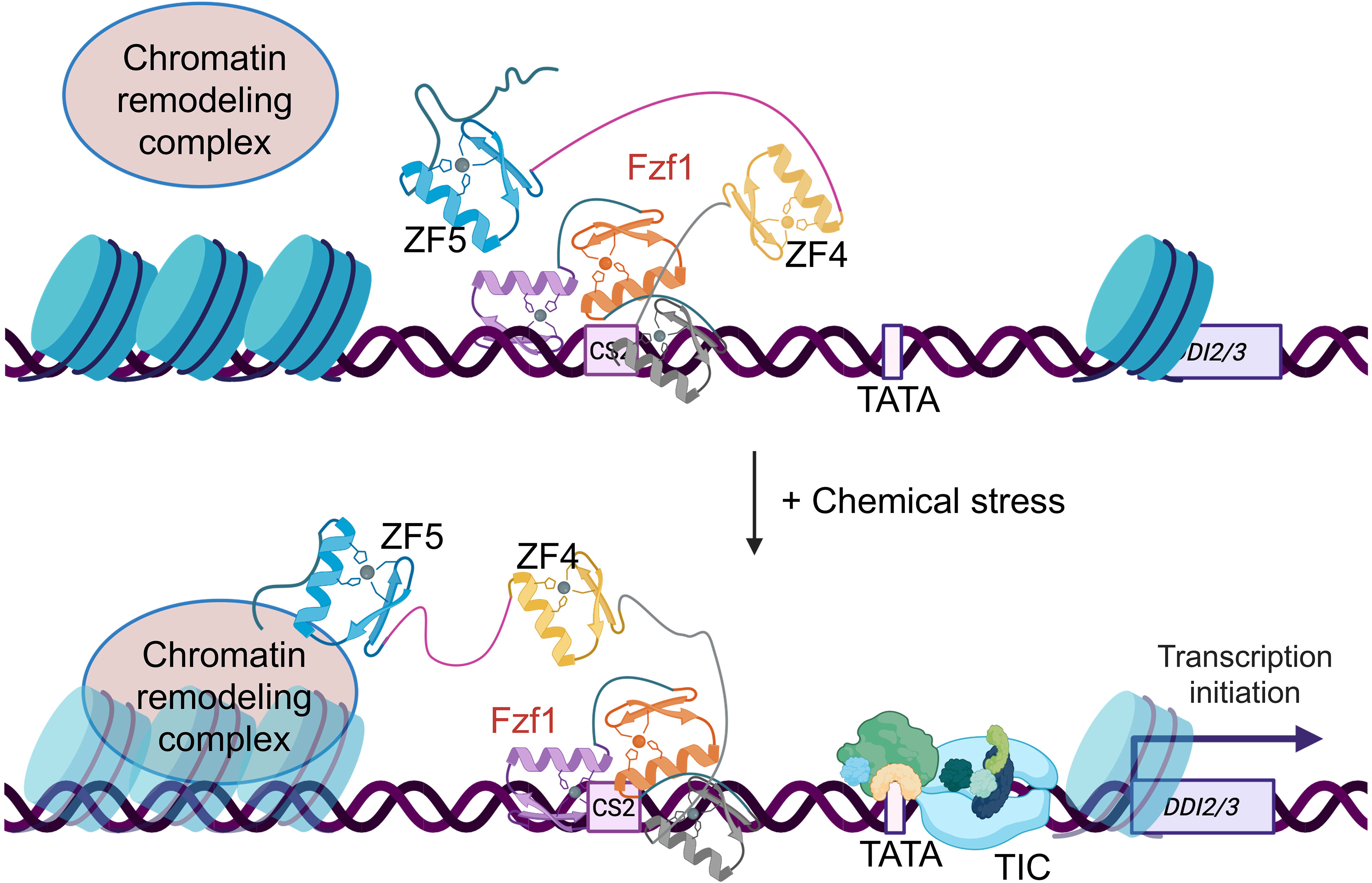
A proposed working model illustrating dual regulation of *DDI2/3* genes by cFzf1 and chromatin remodeling. In the absence of environmental stress, Fzf1 loosely associates with CS2 in an inactive or less active state. Upon exposure to chemical stresses such as CY or MMS, Fzf1 undergoes post-translational modifications (PTMs) and conformational change that relieves the inhibitory effect of ZF4 on the ZF5 activation domain, promoting the interaction with and recruitment of chromatin remodeling factors to the *DDI2/3* promoters, resulting in reduced nucleosome occupancy and increased *DDI2/3* expression. The proposed Fzf1 PTMs, subsequent conformational changes and their roles in the chromatin remodeling have not been experimentally demonstrated. TATA, the TATA box; TIC, transcription initiation complex. Figure was created by using BioRender with academic licence. Created in BioRender. Luo, B. (2026) https://BioRender.com/2jq4gpt.

Although this study revealed the involvement of nucleosome structure in the control of Fzf1-regulated gene expression and dissected the dual requirements of Fzf1 for both nucleosome displacement and target gene activation in the *DDI2/3* promoters in response to certain chemical stresses, it remains unclear how Fzf1 interacts with the nucleosome structure under stress conditions. Pioneer TFs (PTFs) are a subfamily of TFs capable of accessing higher-order chromatin structures (Magnani et al., 2011). PTFs were mainly found and characterized in higher eukaryotes (Zaret and Carroll, 2011). Interestingly, Fzf1 DNA-binding domains show highest primary sequence and structure similarity with mammalian Kif4 (Ma et al., 2026), a multi-functional protein involved in cell differentiation and reprograming (Feinberg et al., 2007; Takahashi and Yamanaka, 2006; Zaehres and Scholer, 2007), is a well-characterized PTF (Soufi et al., 2012; Soufi et al., 2015). In contrast, very few yeast PTFs have been characterized to date (Donovan et al., 2019; Lee et al., 2007b; Mivelaz et al., 2020). A systematic survey of budding yeast TFs with nucleosome-displacing factors (NDFs) revealed that 29 out of 104 TFs contained NDFs, while Fzf1 did not (Yan et al., 2018). However, this study was conducted under untreated conditions, and our MNase-seq results showed that the nucleosome displacement at *DDI2/3* promoters required both CY treatment and functional Fzf1. One possibility is that, upon chemical treatments like CY and MMS, Fzf1 undergoes PTMs to become an NDF. An alternative possibility is that Fzf1 utilizes its ZF5 activation domain (Du et al., 2026) to interact with chromatin remodelers to displace nucleosome (Fig. 9). Future investigation is required to address the above possibilities.

## Materials and methods

### Key Resources Table

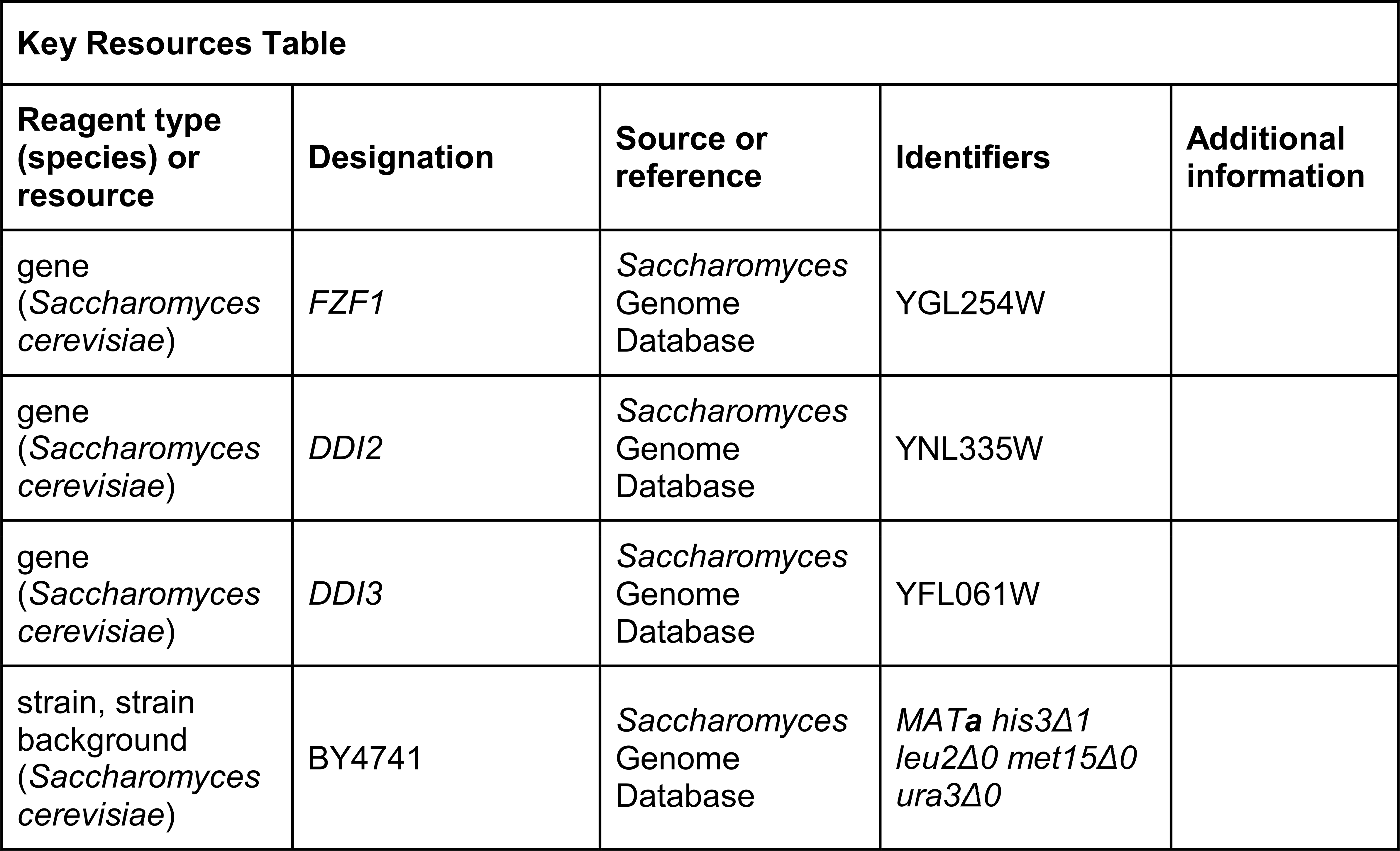
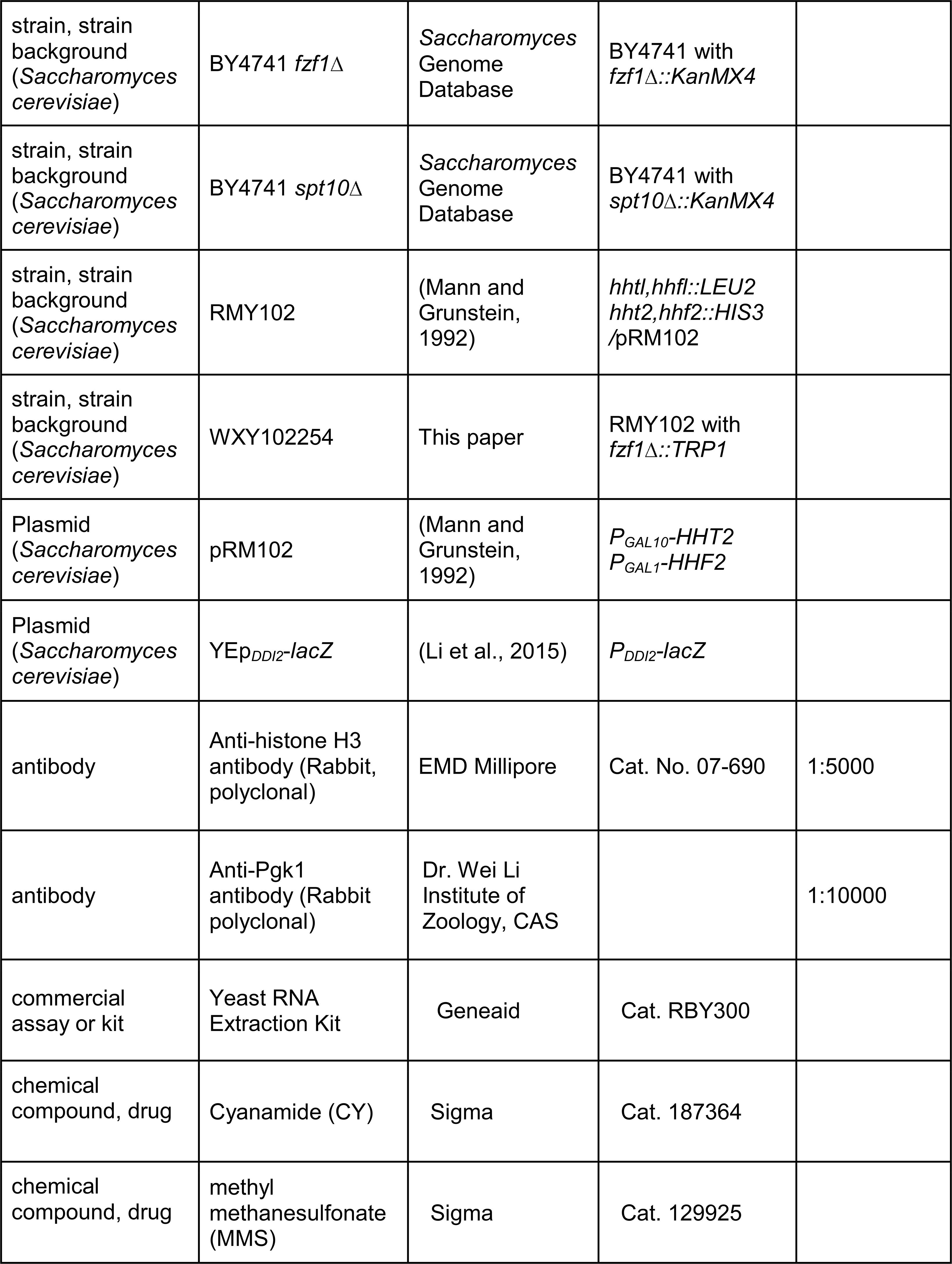
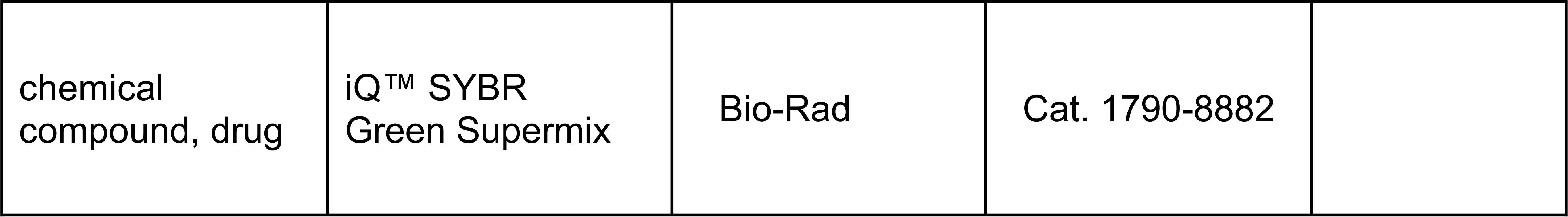

| Key Resources Table |  |  |  |  |
| Reagent type (species) or resource | Designation | Source or reference | Identifiers | Additional information |
|---|---|---|---|---|
| gene (*Saccharomyces cerevisiae*) | *FZF1* | *Saccharomyces* Genome Database | YGL254W |  |
| gene (*Saccharomyces cerevisiae*) | *DDI2* | *Saccharomyces* Genome Database | YNL335W |  |
| gene (*Saccharomyces cerevisiae*) | *DDI3* | *Saccharomyces* Genome Database | YFL061W |  |
| strain, strain background (*Saccharomyces cerevisiae*) | BY4741 | *Saccharomyces* Genome Database | *MAT***a** *his3Δ1 leu2Δ0 met15Δ0 ura3Δ0* |  |
| strain, strain background (*Saccharomyces cerevisiae*) | BY4741 *fzf1*Δ | *Saccharomyces* Genome Database | BY4741 with *fzf1*Δ*::KanMX4* |  |
| strain, strain background (*Saccharomyces cerevisiae*) | BY4741 *spt10*Δ | *Saccharomyces* Genome Database | BY4741 with *spt10*Δ*::KanMX4* |  |
| strain, strain background (*Saccharomyces cerevisiae*) | RMY102 | (Mann and Grunstein, 1992) | *hhtl,hhfl::LEU2 hht2,hhf2::HIS3* /pRM102 |  |
| strain, strain background (*Saccharomyces cerevisiae*) | WXY102254 | This paper | RMY102 with *fzf1*Δ*::TRP1* |  |
| Plasmid (*Saccharomyces cerevisiae*) | pRM102 | (Mann and Grunstein, 1992) | *P<sub>GAL10</sub>-HHT2 P<sub>GAL1</sub>-HHF2* |  |
| Plasmid (*Saccharomyces cerevisiae*) | YEp<sub>*DDI2*</sub>-*lacZ* | (Li et al., 2015) | *P<sub>DDI2</sub>-lacZ* |  |
| antibody | Anti-histone H3 antibody (Rabbit, polyclonal) | EMD Millipore | Cat. No. 07-690 | 1:5000 |
| antibody | Anti-Pgk1 antibody (Rabbit polyclonal) | Dr. Wei Li Institute of Zoology, CAS |  | 1:10000 |
| commercial assay or kit | Yeast RNA Extraction Kit | Geneaid | Cat. RBY300 |  |
| chemical compound, drug | Cyanamide (CY) | Sigma | Cat. 187364 |  |
| chemical compound, drug | methyl methanesulfonate (MMS) | Sigma | Cat. 129925 |  |
| chemical compound, drug | iQ™ SYBR Green Supermix | Bio-Rad | Cat. 1790-8882 |  |

### Yeast strains, cell culture and chemical treatments

Isogenic *ace2*Δ*::KanMX4*, *swi5*Δ*::KanMX4*, *adr1*Δ*::KanMX4*, *ash1*Δ*::KanMX4*, *bas1*Δ*::KanMX 4*, *gcn4*Δ*::KanMX4*, *crz1*Δ*::KanMX4*, *hac1*Δ*::KanMX4*, *mot3*Δ*::KanMX4*, *nrg1*Δ*::KanMX4*, *rgt1* Δ*::KanMX4*, *stb5*Δ*::KanMX4*, *tec1*Δ*::KanMX4* and *spt10*Δ*::kanMX4* strains were derived from BY4741 (*MAT**a** his3*Δ*1 leu2*Δ*0 met15*Δ*0 ura3*Δ*0*) and obtained from the *S. cerevisiae* gene deletion stock. Yeast cells were cultured in yeast extract-peptone-dextrose (YPD) medium or synthetic dextrose (SD) medium supplemented with different nutrients, following standard protocols (Sherman et al., 1983). Strain RMY102, in which both genes encoding histone H3 and H4 are disrupted by *LEU2* and *HIS3* (*hhtl,hhfl::LEU2 hht2,hhf2::HIS3*), and cells were maintained viable by a plasmid pRM102 harboring *HHT2* (H3) and *HHF2* (H4) genes driven by *GAL10* and *GAL1* promoters, respectively (Mann and Grunstein, 1992), and grown in a medium with galactose as the sole carbon source. Strain WXY102254 was created by transforming RMY102 cells with a *fzf1*Δ*::TRP1* cassette, which was produced by PCR amplification using a primer pair H3/H4-FZF1-WF and H3/H4-FZF1-WR (Table Sagainst a template from plasmid YDp-W (Berben et al., 1991). The resulting transformants were confirmed by genomic DNA PCR and sequencing.

For chemical treatments, cells were first grown overnight at 30□°C in YPD or SD selective medium. The overnight cultures were then diluted into fresh medium at 600□nm (OD□□□_nm_) of 0.2–0.3 and allowed to grow for approximately 2 hours until reaching exponential phase. At this point, cells were treated with desired concentrations of test chemicals, including CY or MMS for 2 hours. Control cells were incubated under the same conditions without chemical treatment.

### Construction of plasmid YEp*_DDI2_*-*lacZ* and promoter deletions

Plasmid YEp*_DDI2_*-*lacZ* was constructed by cloning a 1.17-kb *Hin*dIII-*Pst*I fragment containing 709-bp *DDI2* promoter and a 461-bp coding sequences into plasmid YEp356R (Myers et al., 1986) as previously described (Li et al., 2015).

To make *DDI2* promoter deletions from plasmid YEp*_DDI2_*-*lacZ*, the available unique restriction sites, including *Spe*I (−453), *Xba*I (−358) and *Bgl*II (−229) (Figure S1), were utilized. In addition, promoter deletions up to -190, -90, -70, -50 and -30 were made by using primers as shown in Supplementary Table S2 for the PCR reaction. To delete the 6-bp repeat sequences 5′-AAAAGA-3′ from the *DDI2* promoter, D1, D2, D3 and D4 were sequentially deleted from plasmid YEp*_DDI2_*-*lacZ* by using forward and reverse primer pairs as listed in Table S2.

### Yeast cell transformation

Yeast cells were transformed with plasmids using a modified lithium acetate method (Ito et al., 1983) as described (Gietz and Woods, 2006).

### Yeast reporter gene assay

To investigate downstream gene expressions under the control of various *DDI2/3* promoter truncations, a series of truncated YEp*_DDI2_*-*lacZ* plasmids were transformed into the wild-type yeast strain BY4741. Transformants selected on SD-Ura plates were used to inoculate liquid SD-Ura medium. Overnight cultures were diluted into fresh selective medium to an OD□□□_nm_ of 0.2 and incubated for an additional 2 hours to reach exponential phase, followed by chemical treatments as described above. β-Galactosidase (β-Gal) activity was measured using the standard assay with ortho-nitrophenyl-β-galactoside (ONPG) as the substrate, and the enzymatic activity was quantified by measuring absorbance at 420 nm (OD□□□_nm_), as previously described (Li et al., 2015).

### Yeast RNA extraction and quantitative reverse transcription PCR (RT-qPCR)

Following chemical treatments, yeast cells were harvested by centrifugation and subjected to zymolyase (Amsbio, Cat. 120491-1) treatment at 200 U per 5 × 10□ cells for 1 hour to lyse the cells. Total RNA was extracted using the Yeast RNA Extraction Kit (Geneaid, Cat. RBY300). The purified RNA was reverse transcribed into cDNA for long-term storage and subsequent experiments. Gene expression levels were quantified by quantitative PCR using iQ™ SYBR Green Supermix (Bio-Rad, Cat. 170-8882). Relative gene expression was calculated using the 2^□ΔΔCT^ method (Schmittgen and Livak, 2008), with normalization to the internal reference gene *UBC6* as previously validated (Lin et al., 2023).

### Western blot analysis

Yeast cellular histone H3 protein level was monitored by western blot analysis. Total protein extracts were prepared using the glass bead lysis method, as previously described (Ball and Xiao, 2014). Histone H3 was detected using an anti-H3 antibody (EMD Millipore, Cat. No. 07-690) at a 1:5000 dilution. An anti-Pgk1 polyclonal antibody, kindly provided by Dr. Wei Li (Institute of Zoology, Chinese Academy of Sciences), was used as an internal loading control. The band intensities were measured, calculated and compared using Adobe Photoshop.

### Micrococcal nuclease sequencing (MNase-seq) assay

MNase-seq was performed as described previously (McKnight et al., 2021). BY4741 wild-type and its isogenic *fzf1*Δ*::KanMX4* mutant cells were grown to log phase and treated with 20 mM CY for 2 hours. Cells were crosslinked with 1% formaldehyde for 15 minutes and quenched by glycine to a final concentration of 300 mM for 5 minutes. Pellets were washed three times with cold 10 mM Tris-Cl pH 7.5, 100 mM NaCl and then flash frozen. Cells were resuspended in spheroplast buffer, and 10% cross-linked *Schizosaccharomyces pombe* cells were added to each sample as a spike-in. Cells were treated with 2 mg of zymolyase for 15 mins at 25°C with rotation, then digested with 60 units of micrococcal nuclease (MNase) and 30 units of ExoIII for 11 minutes at 37°C. Residual proteins and RNAs were removed, and DNA was purified with a QIAGEN MinElute PCR Purification Kit. Libraries were prepared with NEB DNA Ultra II and sequenced on an Illumina NextSeq.

Adapter sequences were removed from paired end FASTQ files using cutadapt (version 4.9), before aligning to the sacCer3 genome using BWA (version 0.7.17-r1188). Samtools view (version 1.21) was used to retain uniquely mapping and properly paired reads with a Mapping Quality (MAPQ) score of 10 or higher. Note the due to sequence similarity between *DDI2* and *DDI3*, approximately 55% of reads mapping to these loci were discarded by this filtering. BAM files were converted to BED files using awk (version 20200816), and coverage tracks, representing reads per genome coverage (RPGC), were calculated using the Java Genomics Toolkit (https://github.com/timpalpant/java-genomics-toolkit) scripts, ngs.BaseAlignCounts and wigmath.Scale. Read coverage over nucleosomes mapping upstream of TSSs (Weiner et al., 2015) was calculated using the Java Genomics Toolkit ngs.IntervalStats tool. Jitter plots were generated using R packages dplyr (version 1.1.4), tidyr (version 1.3.1), and ggplot2 (4.0.0) and heatmaps were created using the Deeptools computeMatrix and plotHeatmap tools (Ramirez et al., 2014) aligning 5,542 protein-coding genes by the TSS. Data generated for this manuscript were deposited in the NCBI Gene Expression Omnibus under the accession code “GSE327798”

### Statistical analyses

Statistical analyses were performed using GraphPad Prism (ver. 9.5.0). Comparisons between datasets were analyzed using a two-way analysis of variance (two-way ANOVA) to evaluate the effects of multiple independent variables. Multiple comparisons tests were applied following ANOVA to determine statistical differences between individual groups. A *P* value < 0.05 was considered statistically significant.

## Supplementary data

Contains two tables and six figures.

## Data availability

MNase-seq data generated for this manuscript were deposited in the NCBI Gene Expression Omnibus under the accession code “GSE327798”, and can be accessed through the following website: https://genome.ucsc.edu/s/ljhowe%40mail.ubc.ca/Xiao

## Acknowledgements

The authors wish to thank Dr. Michael Grunstein for the yeast strain RMY102 and Dr. Wei Li for the anti-Pgk1 antibody. This work was supported by the Natural Sciences and Engineering Research Council of Canada Discovery Grants RGPIN-2025-06661 to WX and RGPIN-2024-06053 to LJH.

## Additional information

### Author Contributions

Conceptualization, YD, AL and WX; experiments, YD, AL and JB; data analysis, YD, AL, LJH and WX; writing – original draft preparation, YD; review and revision, WX and LJH.

Supervision, WX and LJH; funding requisition, WX and LJH. All authors have read and agreed to the published version of the manuscript.

