## Supplemental tables and figures for "Dual regulation of chemical stress-induced *DDI2/3* expression by a transcription factor Fzf1 and nucleosome in *Saccharomyces cerevisiae*"

### Supplementary data

**Table S1.** Comparison of effects of histone depletion on Fzf1-regulated gene expression from this study by RT-qPCR and a previous microarray report (Wyrick et al., 1999).

| | **Wild type** | ***fzf1*Δ** | **Wyrick, 1999** |
|---|---|---|---|
| ***DDI2/3*** | **29.4 (4.8)** | **11.2 (3.62)** | **68.5/305.8** |
| ***SSU1*** | **0.6 (0.13)** | **1.1 (0.37)** | **N/A** |
| ***YHB1*** | **0.1 (0.02)** | **0.2 (0.21)** | **-5.1** |
| ***YNR064C*** | **7.7 (1.59)** | **5.9 (2.45)** | **20.1** |

**Table S2**. Oligonucleotide sequences used in this study

| **Name** | **Sequence (5'-3')** |
|---|---|
| DDI2/3-F | GGAAAATCCAAGCTTTCAAG |
| DDI2/3-R | GCCGCGGCCGCCCTCATTGAAACTTACCT |
| UASΔ F | CGAATCAAGCTTACAAAATACTCCACCGCA |
| UASΔ R | CAATTCTGCAGAATGAGCCCCAAGGTGGT |
| ΔN-90 F | CGACGTAAGCTTTATATAAATGGGTTAGTT |
| ΔN-70 F | AGCCACAAGCTTCTTCTTCTTCTGTTAACA |
| ΔN-50 F | CGAACCAAGCTTAAGTTGCTTCGTACTGTT |
| ΔN-30 F | ATCATGAAGCTTTTGCCTTGCTTCTCTTCA |
| D1 F | GATTTTTCTGACTCCAAACAGAGCCCTGCGATAGTT |
| D1 R | AACTATCGCAGGGCTCTGTTTGGAGTCAGAAAAATC |
| D2 F | GACTGAATCCCTCAAGAAAATTGCCTAACTGATACTGG |
| D2 R | CCAGTATCAGTTAGGCAATTTTCTTGAGGGATTCAGTC |
| D3 F | CTAACTGATACTGGTTTAGAAAGATGTCAAATATGC |
| D3 R | GCATATTTGACATCTTTCTAAACCAGTATCAGTTAG |
| D4 F | CGGCCCCGAAACAAATGTCCGCAATCTTATTAAAGTGC |
| D4 R | GCACTTTAATAAGATTGCGGACATTTGTTTCGGGGCCG |
| H3/H4-*FZF1*-WF | AAGAAGATGAAAATGACAACTTTTACGCTGGTGTGCACAA GTGGTACCAGAATACGTGGCCACCAAAACAGAATTCCCGG GGATCCGGTG |
| H3/H4-*FZF1*-WR | CTCTTGCCTTTTCTTTGTTTCATAGTTCGAATCACATGAGTA GAGGACGGAAATTGCTCTTCTATGGCGTCTGCAGGTCGACG GATCCGG |

### Supplementary figure elegends

**Figure S1. DNA sequences at *DDI2* and *DDI3* promoters**.

The *DDI2* promoter sequence is shown with nucleotide variations in *DDI3* indicated in orange letters. Restriction sites used in this study for molecular cloning and truncations are indicated. The 5'-AAAAGA-3' repeats are underlined and indicated in green. The CS2 Fzf1 recognition sequence is indicated in magenta, and the TATA box sequence is indicated in blue. The bold ATG at +1 is the translation start codon in *DDI2* and *DDI3*.

**Figure S2. Effects of the 6-bp 5'-AAAAGA-3' repeat sequence deletions on the $P_{DDI2}$-*lacZ* expression.**

(A) Illustration of sequential 6-bp repeat sequence deletions. The deletion constructs were designated Δ1 through Δ1234. Solid green bars represent the 6-bp repeat sequences and the dashed bars represent deletion of the 6-bp sequences. See Fig. S1 for the exact location of these repeats. (B) Fold inductions of the $P_{DDI2}$-*lacZ* reporter gene under the 20 mM CY and 2-hr treatment conditions in BY4741 cells harboring the indicated 6-bp deletion plasmids. Errors represent standard deviations from at least three independent experiments.

**Figure S3. Effects of transcription repressor gene deletions on the $P_{DDI2}$-*lacZ* expression**.

(A) Putative binding sites for 15 transcription repressors within the defined 480-nt URS region (nt -709 to -229 bp) identified through the SGD website. (B) β-Gal activities to measure $P_{DDI2}$-*lacZ* expression in the 13 indicated BY4741 transcription repressor deletion strains. All $YEp_{DDI2}$-*lacZ* transformed cells were treated with 20 mM CY for 2 hours. Errors represent standard deviations from at least three independent experiments.

**Figure S4. RMY102 cell growth curves in liquid YPGal (green) and YPD (red) media incubated at 30 °C with shaking.**

Samples were taken from cultures at indicated periods up to 7 hours and the $OD_{600\ nm}$ were determined by spectrophotometry and converted to the cell density with a standard curve. Errors represent standard deviations from at least three independent experiments.

**Figure S5. Western blot analysis of cellular histone H3 levels in histone depleted RMY102 cells under CY and MMS treatment conditions.**

RMY102 cells were grown in the YPD medium under H3/4 depletion conditions for 4 hours and then treated with 20 mM CY or 0.1% MMS for 2 hours. The whole-cell lysates were subjected to immunoblotting using anti-H3 antibodies (upper panel). Anti-Pgk1 (lower panel) blotting served as an internal control. Numbers underneath the anti-H3 blot indicate histone H3 levels relative to cells grown in YPD without chemical treatments.

**Figure S6. Quality controls of the MNase-seq assay.**

(A) Purified DNA samples used for the MNase-seq analyzed on a High Sensitivity D1000 ScreenTape using an Agilent TapeStation. (B) Metagene plots and heat maps representing MNase-seq read counts normalized to the genomic mean at 5542 genes aligned by their TSS (Chereji et al., 2018). Heatmaps are ordered by the width of the nucleosome depleted region.

**Figure S7. Nucleosome occupancy at selected loci under various experimental conditions**.

BY4741 (WT) and BY4741 *fzf1*Δ cells with or without 20 mM CY treatment for 2 hours were subjected to MNase-seq and genome browser tracks of sequence coverage (RPGC) over *SSU1* (A), *YHB1* (B), *YNR064C* (C) are shown.

```
               HindIII
-720 TGGAAAATCCAAGCTTTCAAGATGGGTAACTGTTATTCAAAGGATCCTCTAAGATAAAAC
DDI3                         C

-660 ACAGATCGACAGATCCGAGAGTTGGCTTCTGTGCCTTGGGCTCAAATTCCTTTCCCACCT
DDI3         G

-600 CATGCAAATTGATTTTTCTGACTCCAAAAAAAGACAGAGCCCTGCGATAGTTCCCGAATG
DDI3    T                                A

-540 TTGTAACATCAAAGCCAAGCACTCCTTTATAGAAGTCGCATGAACGTTGAATACTAGCAG

                               SpeI
-480 CTGGTGAAACTACAGGGTCTAAACTAACTAGTATCCATATCTTTTTGAGAGCATTGAAAG
DDI3                                          G

-420 TATACGGAGTACAAGCTGGGTTAGAAGGAATTTTTATCTTAACAGCAATGAAAATCAACT

      XbaI
-360 TTCTAGACTGAATCCCTCAAGAAAATTGCAAAAGACTAACTGATACTGGTTTAAAAGAGA
DDI3                                         C

-300 AAGATGTCAAATATGCGGAGTTATACCATCAAACAACTTTGGACGGCCCCGAAACAAATG

               BglII                      CS2
-240 TCCGCAAAAAAGATCTTATTAAAGTGCATGGACACTATCATTTCTATAATACAAAATACT

-180 CCACCGCACAATAGTTTGTCGGGAAGTCATCAATCAATCTTGTACGAGCTTTACAAATAA

                                    TATA box
-120 CTTTTTAGGATCGGTCCCCTCATAAAATTATATATAAATGGGTTAGTTTCCTTCTTCTTC

 -60 TGTTAACATGAAGTTGCTTCGTACTGTTTTTTGCCTTGCTTCTCTTCAAAAGAATCAATC
DDI3                                         C

   1 ATGTCACAGTACGGATTTGTAAGAGTTCCTAGAGAGGTAGAAAAGGCCATTCCAGTGGTG
```

Figure S1

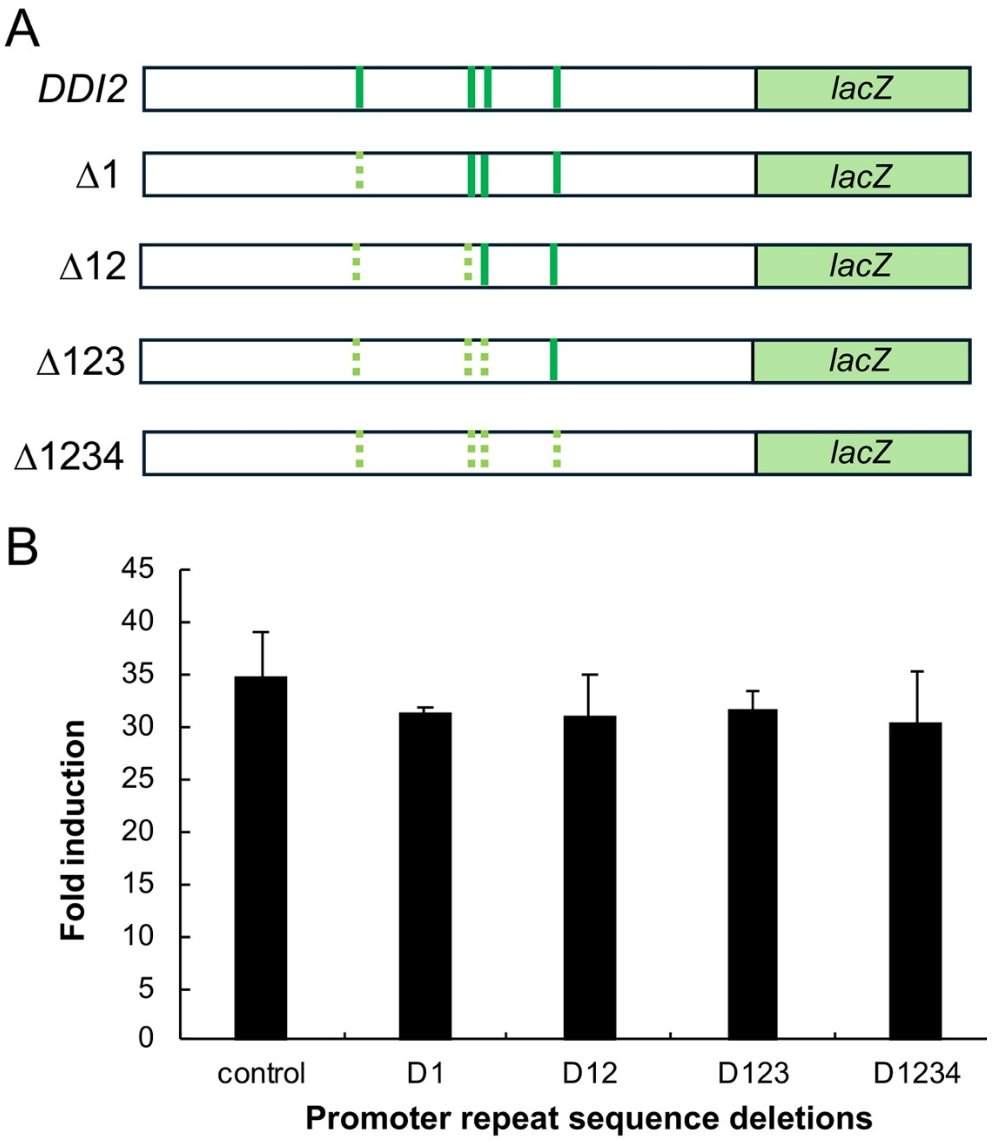
A
DDI2
lacZ
Δ1
lacZ
Δ12
lacZ
Δ123
lacZ
Δ1234
lacZ
B
45
40
35
30
25
20
15
10
5
0
Fold induction
control
D1
D12
D123
D1234
Promoter repeat sequence deletions

Figure S2

A

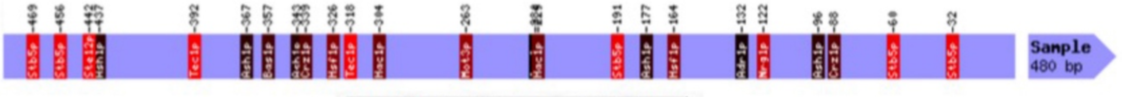


| Transcription Factor | Consensus | Position | Strand |
|---|---|---|---|
| Target Sequence: Sample (size 480) Back to top | | | |
| Ace2p, Swi5p | ACCAGC | -250 | R |
| Adr1p | TTGGRG | -348 | R |
| Ash1p | YTGAT | -367 | F |
| Ash1p | YTGAT | -96 | F |
| Ash1p | YTGAT | -43 | R |
| Ash1p | YTGAT | -137 | R |
| Ash1p | YTGAT | -303 | R |
| Bas1p, Gcn4p | TGACTC | -357 | F |
| Crz1p | GAGCCC | -339 | F |
| Crz1p | GAGCCC | -392 | R |
| Hac1p | CCAGC | -176 | R |
| Hac1p | CCAGC | -251 | R |
| Hsf1p | NTTCNNGAAN | -326 | F |
| Hsf1p | NTTCNNGAAN | -316 | R |
| Mot3p | ATGGAT | -217 | R |
| Mot3p | AAGGWT | -456 | F |
| Nrg1p | CCCTC | -122 | F |
| Rgt1p | CGGANNA | -191 | F |
| Stb5p | CGGNS | -191 | F |
| Stb5p | CGGNS | -60 | F |
| Stb5p | CGGNS | -32 | F |
| Stb5p | CGGNS | -11 | R |
| Stb5p | CGGNS | -24 | R |
| Ste12p | ATAAAACA | -442 | F |
| Tec1p | RMATTCYY | -392 | F |
| Tec1p | RMATTCYY | -162 | R |

B

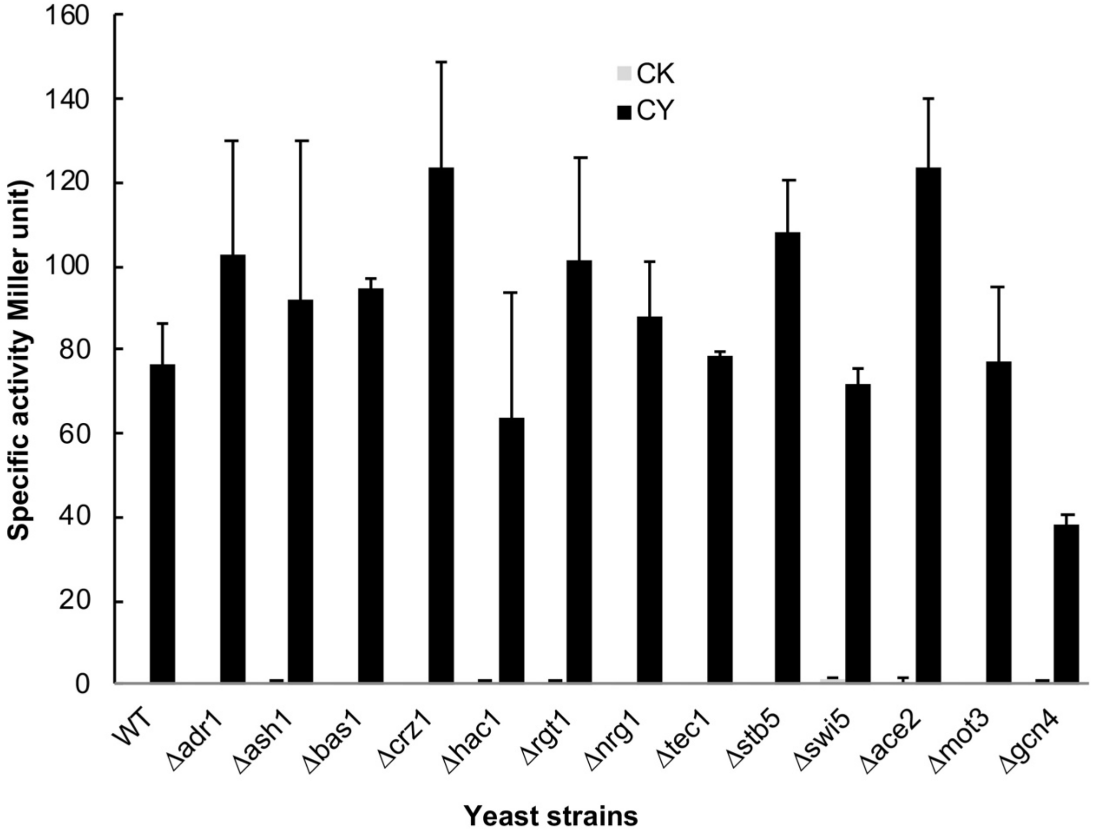


Figure S3

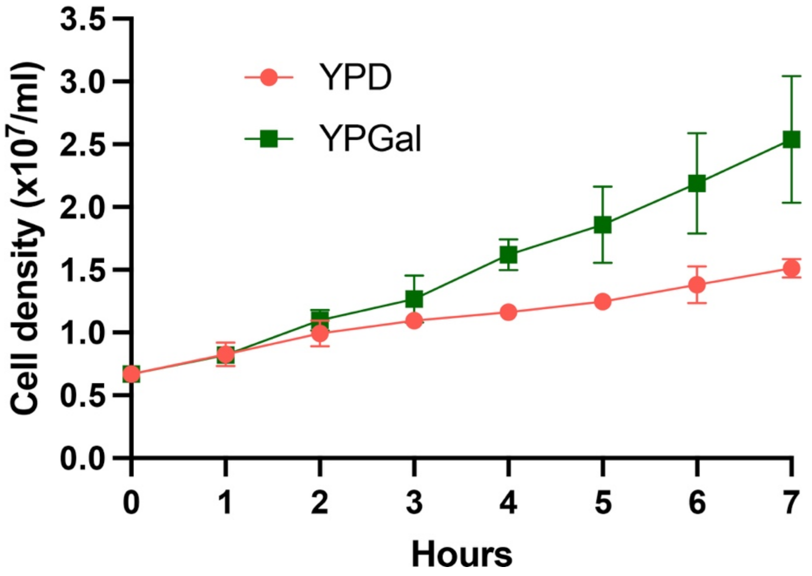
YPD
YPGal
Cell density (x10^7/ml)
3.5
3.0
2.5
2.0
1.5
1.0
0.5
0.0
0
1
2
3
4
5
6
7
Hours

Figure S4

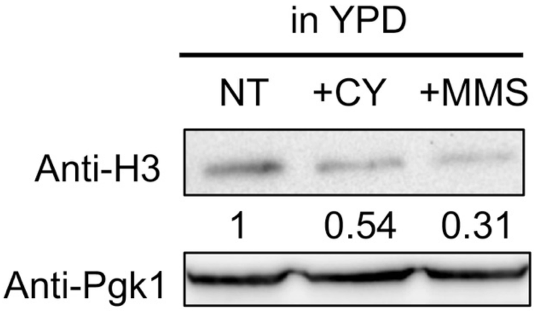


Figure S5

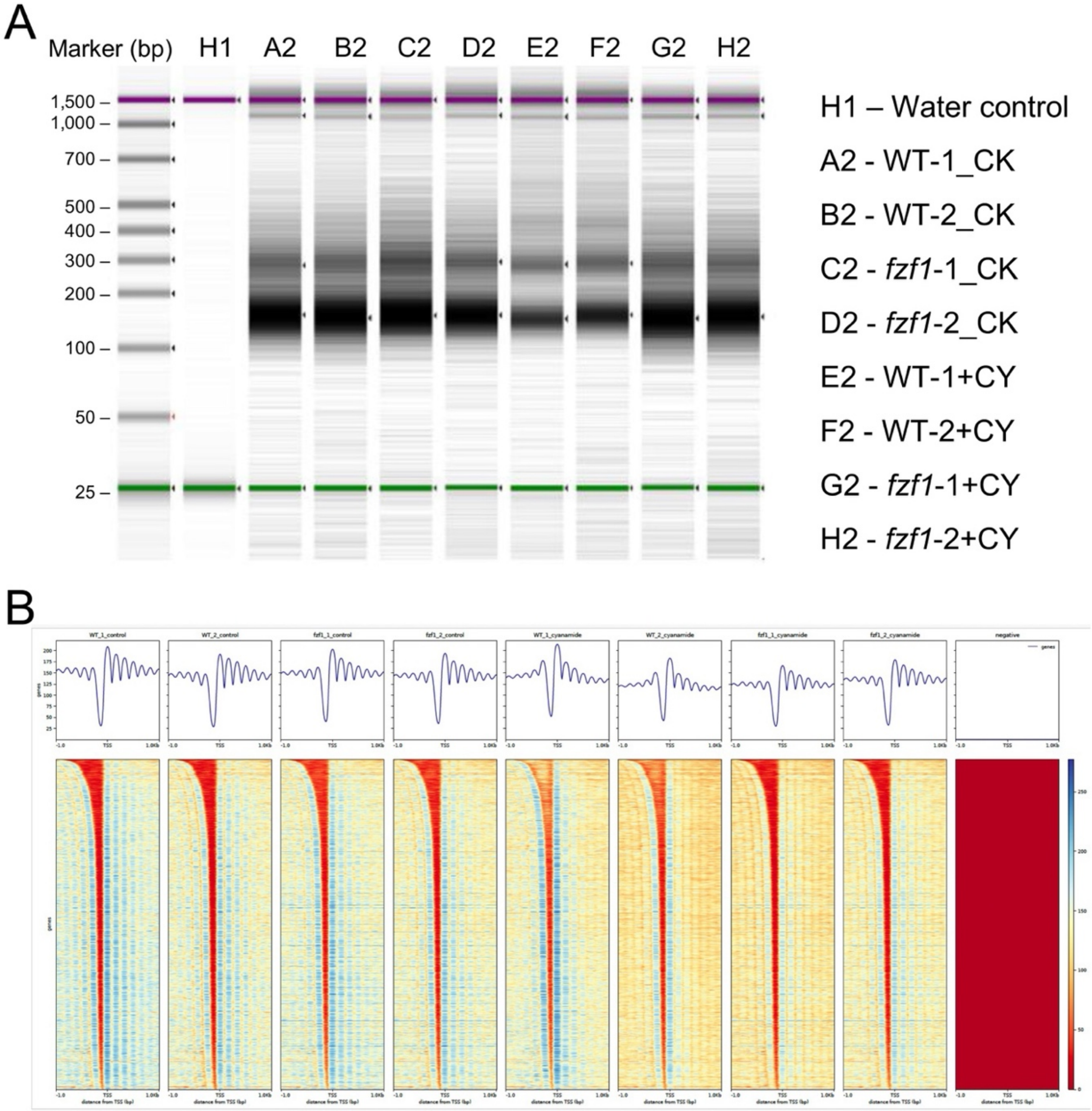
A
Marker (bp)
H1
A2
B2
C2
D2
E2
F2
G2
H2
1,500 –
1,000 –
700 –
500 –
400 –
300 –
200 –
100 –
50 –
25 –
H1 – Water control
A2 - WT-1_CK
B2 - WT-2_CK
C2 - fzf1-1_CK
D2 - fzf1-2_CK
E2 - WT-1+CY
F2 - WT-2+CY
G2 - fzf1-1+CY
H2 - fzf1-2+CY
B
WT_1_control
WT_2_control
fzf1_1_control
fzf1_2_control
WT_1_cyanamide
WT_2_cyanamide
fzf1_1_cyanamide
fzf1_2_cyanamide
negative
genes
TSS
-1.0
1.0Kb

Figure S6

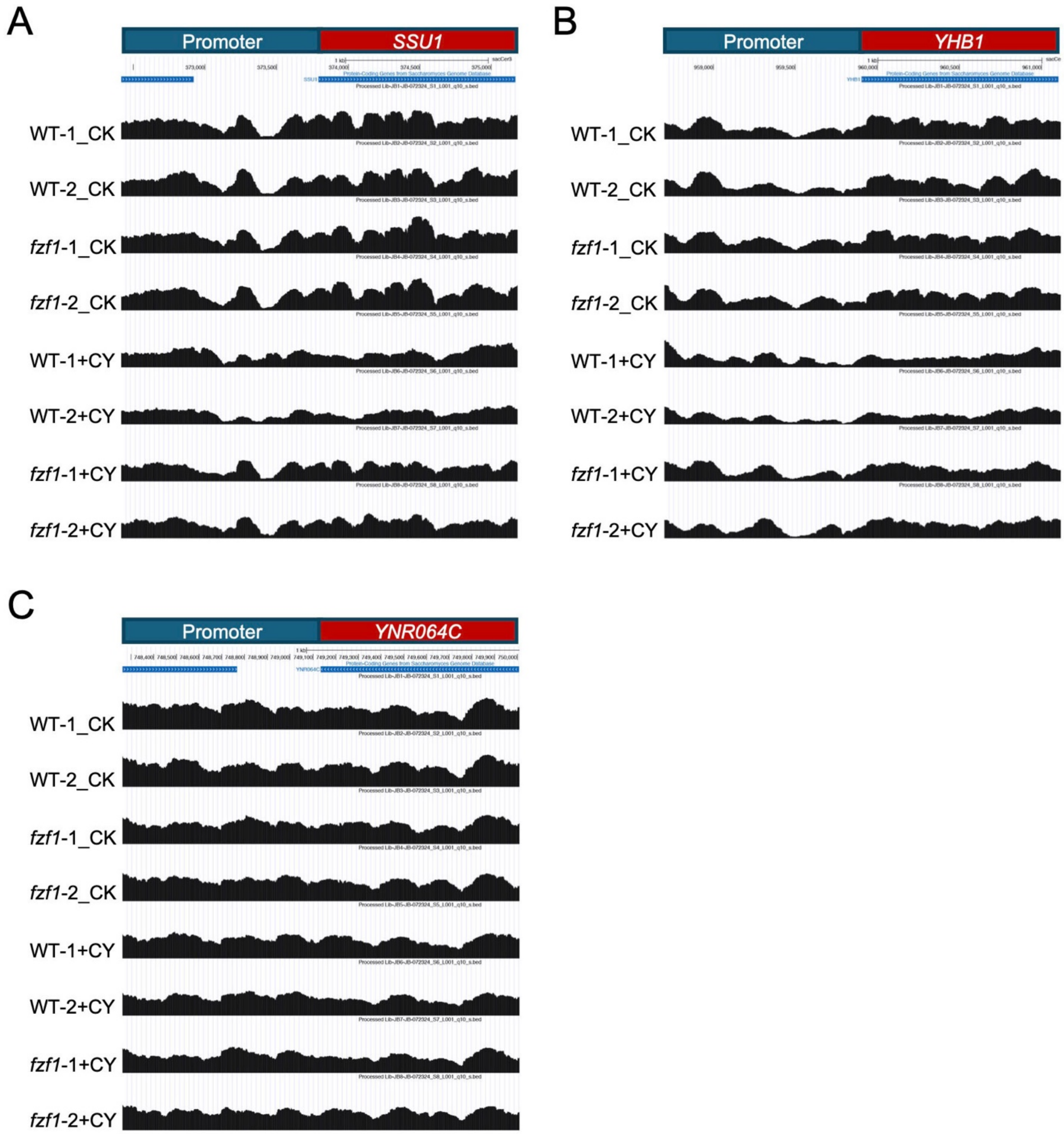
A
Promoter
SSU1
WT-1_CK
WT-2_CK
fzf1-1_CK
fzf1-2_CK
WT-1+CY
WT-2+CY
fzf1-1+CY
fzf1-2+CY
B
Promoter
YHB1
WT-1_CK
WT-2_CK
fzf1-1_CK
fzf1-2_CK
WT-1+CY
WT-2+CY
fzf1-1+CY
fzf1-2+CY
C
Promoter
YNR064C
WT-1_CK
WT-2_CK
fzf1-1_CK
fzf1-2_CK
WT-1+CY
WT-2+CY
fzf1-1+CY
fzf1-2+CY

Figure S7
